# Decoding differential gene expression

**DOI:** 10.1101/2020.01.10.894238

**Authors:** Shinya Tasaki, Chris Gaiteri, Sara Mostafavi, Yanling Wang

## Abstract

Identifying the molecular mechanisms that control differential gene expression (DE) is a major goal of basic and disease biology. Combining the strengths of systems biology and deep learning in a model called *DEcode*, we are able to predict DE more accurately than traditional sequence-based methods, which do not utilize systems biology data. To determine the biological origins of this accuracy, we identify the most predictive regulators and types of regulatory interactions in DEcode, contrasting their roles across many human tissues. Diverse systems biology, ontological and disease-related assessments all point to the predominant influence of post-translational RNA-binding factors on DE. Through the combinatorial gene regulation that is captured in DEcode, it is even possible to predict relatively subtle person-to-person variation in gene expression. We demonstrate the broad applicability of these clinically-relevant predictions by predicting drivers of aging throughout the human lifespan, gene coexpression relationships on a genome-wide scale, and frequent DE in diverse conditions. Researchers can freely access DEcode to utilize genomic big data in identifying influential molecular mechanisms for any human expression data - www.differentialexpression.org.

## Introduction

While all human cells share DNA sequences, gene regulation differs among cell types and developmental stages, and in response to environmental cues and stimuli. Accordingly, when gene expression is not properly regulated, cellular homeostasis can be perturbed, often affecting cell function and leading to disease^1^. These distinctions between cell states are observed as differential expression (DE) of gene transcripts. DE have been cataloged for tens of thousands of gene expression datasets, in the context of distinctions between species, organs, and conditions. Despite the important and pervasive nature of DE, it has been challenging to shift from these observations towards a coherent understanding of the underlying generative processes that would essentially decode DE– a transition which is essential for progress in basic and disease biology. We address this gap by exploiting novel computational and systems biology approaches to develop a predictive model of DE based on genome-wide regulatory interaction data. Utilizing diverse genomic datasets, we identify a complex, yet strikingly consistent set of principles that control DE. This model of differential expression, called DEcode, can be applied to the majority of current and future gene expression data, to accelerate basic and disease biology, by identifying the origins of DE in each experiment.

Diverse molecular interactions have been shown to generate DE, and jointly regulate gene expression at the transcriptional and post-transcriptional levels. Major classes of gene regulatory interactions have been cataloged at the genomic scale, including transcription factor (TF)-promoter interactions^2^, protein-RNA interactions^3^, RNA-RNA interactions^4^, chromatin interactions^5^, and epigenetic modifications on DNAs^6^, histones, and RNAs^7^. Statistical models of gene expression can help fulfill the purpose of these resources in describing the origins of gene regulation and DE^1^. However, such raw data resources have outpaced model development, likely due to the challenge of uniting diverse molecular data into a single accurate model.

Predicting DE on the basis of gene regulatory interactions is one initial approach to understanding its origins. Among many possible statistical approaches to predicting DE, deep learning (DL) blends diverse data sources in a way that approximates the convergence of regulatory interactions. Indeed, DL has been applied to genomic research^8, 9^ including RNA splicing^10^, genomic variant functions^11^, and RNA/DNA binding^12^. However, accurate prediction is only one component of understanding DE; additional genomic and systems biology analysis are helpful in understanding how predictions are fueled by existing molecular concepts, mechanisms, and classes.

To decode the basis of DE in terms of molecular regulatory interactions, we first learn to predict it with a high degree of accuracy, using a DL model we call “DEcode”. This model combines several types of gene regulatory interactions and allows us to prioritize the main systems and molecules that influence DE on a tissue-specific basis. We further establish likely molecular mechanisms for this gene regulation and validate the influence of the predicted strongest regulators. In parallel, we predict the origin of person-to-person DE, which is the major component of experimental and clinical studies. These particularly challenging predictions are validated on a genome-wide scale, as we identify key drivers of coexpression, and also drivers for phenotype-associated differential expression. These tests and applications indicate DEcode can combine multiple recent data sources, to extract regulators for arbitrary human DE signatures.

## Results

### Promoter and RNA features predict differential expression across human tissues

The overarching goal of this study is to accurately predict gene expression as a function of molecular interactions. These results should be tissue-specific, but also highlight major regulatory principles across tissues, and ideally have sufficient accuracy to predict the relatively small expression changes observed between individual humans. To accomplish this, we utilized deep convolutional neural networks in a system called DEcode that can predict inter-tissue variations and inter-person variations in gene expression levels from promoter and mRNA features (**Figure 1**). The promoter features included: the genomic locations of binding sites of 762 TFs and the mRNA features encompassed the locations of binding sites of 171 RNA-binding proteins (RNABPs) and 213 miRNAs in each mRNA (**Table S1**). DEcode takes the promoter features and the mRNA features for each gene as inputs and outputs its expression levels under various conditions. We note that the prediction is based on only the presence or absence of known binding sites, and other information such as gene expression levels of TFs, RNABPs, and miRNAs is not utilized. First, we applied the DEcode framework to tissue-specific human transcriptomes of 27,428 genes and 79,647 transcripts measured in the GTEX consortium^13^ to predict log-fold changes across 53 tissues against the median log-TPM (transcripts per million) of all tissues, as well as the median log-TPM of all tissues with a multi-task learning architecture. To ensure rigorous model testing, we excluded all genes or transcripts coded in chromosome 1 from the training data and used them as the testing data for evaluating the performances of DEcode models. This procedure prevents information leaking from intra-chromosomal interactions and potential overlaps of regulatory regions (details of model construction in **Figure S1**). The predicted median TPM levels showed high consistency with the actual observations for both gene-level (Spearman’s rho = 0.81) and transcript-level (Spearman’s rho = 0.62) (**Figure 2A**). Moreover, the model predicted the differential transcript usage within the same gene (Spearman’s rho = 0.44) (**Figure 2A**). The DEcode models also predicted the differential expression profiles across 53 tissues for both gene (mean Spearman’s rho = 0.34), transcript (mean Spearman’s rho = 0.32), and transcript-usage levels (mean Spearman’s rho = 0.16) (**Figure 2B**). The predicted gene expression for the testing genes was indeed tissue-specific, as they showed less correspondence with the expression profiles from alternate tissues (**Figure 2C**).

**Figure 1.**
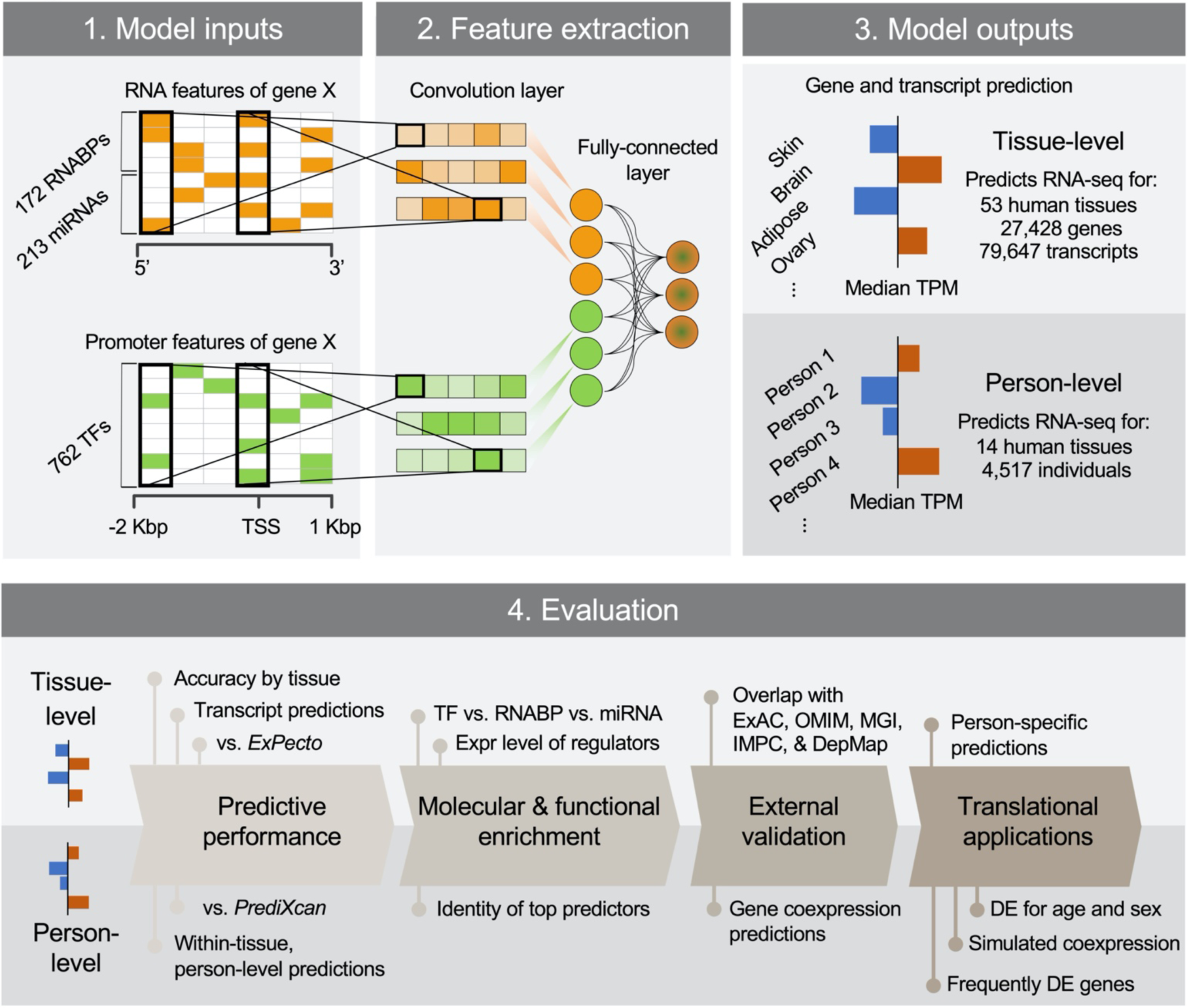
Overview of building and evaluating the DEcode transcriptome prediction model.

**Figure 2.**
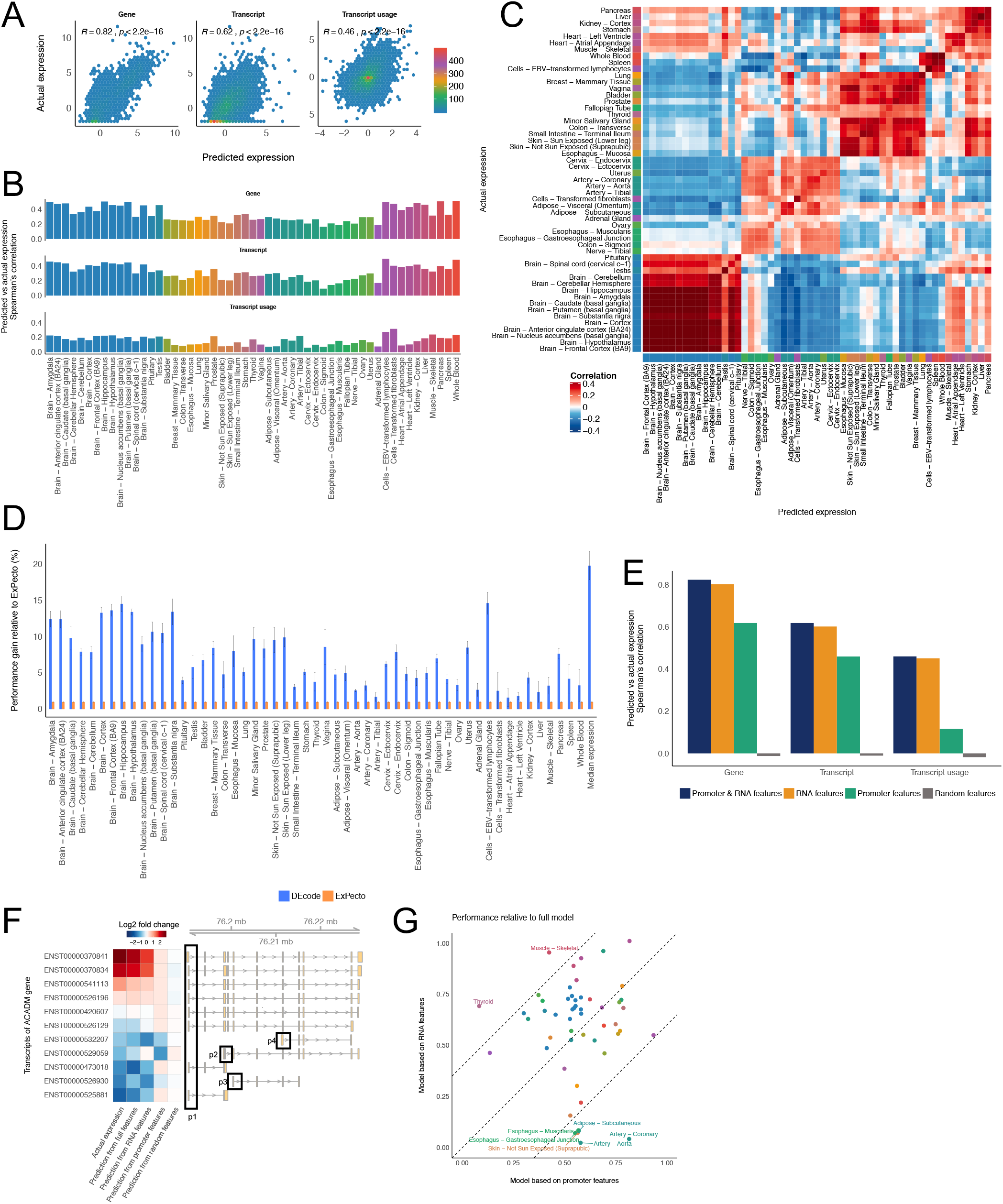
Performance of the tissue-level models. (**A**) Prediction performances on the median absolute expression levels across tissues. The predicted the log2-TPM values for 2,705 genes or 7,631 transcripts coded on chromosome 1 were compared with the actual median log2-TPMs across 53 tissues using Spearman’s rank correlation. The transcript usage within each gene was computed by subtracting the mean log2-TPM from log2-TPM of transcripts in each gene for 1,485 genes that had multiple transcripts. (**B**) Prediction performances on the tissue-specific expression profiles. The predicted fold changes relative to the median of all tissues for 2,705 genes or 7,631 transcripts coded on chromosome 1 were compared with the actual fold changes in each tissue using Spearman’s rank correlation. The differences in the transcript usage within each across tissues were computed for 1,485 genes that had multiple transcripts. The color of the bar indicated the tissue groups based on the similarity of gene expression profiles. (**C**) The heatmap showing pairwise correlations between the predicted and the actual tissue-specific expression profiles of 53 tissues for the testing genes. (**D**) Performance comparison of DEcode with ExPecto. The root-mean-square errors (RMSE) of DEcode models for expression-levels of 714 genes coded on chromosome 8 was compared with those of ExPecto. Each method was executed 10 times. The median RMSE of the 10 runs was displayed as a bar plot and the error bar represents median absolute deviation. (**E**) The predictive performances of the models trained with a different set of features. (**F**) The comparison of the expression levels for ACADM transcripts predicted by the models trained with different feature sets. (**G**) The predictive performances on the tissue-specific gene expression profiles of the testing data relative to the model trained with a full set of features.

To provide context for the statistical performance of DEcode, we contrast it to a high-performing method called ExPecto^11^, as it was designed to predict GTEX gene expression from epigenetic states, estimated from promoter sequences via DL. We built 10 models for each method using the same genes for training, validation, and testing to predict gene expression in the 53 tissues. In this comparison, DEcode showed an average of 7.2% improvement in root mean square error over ExPecto (**Figure 2D**) which translates into an average correlation coefficient with actual gene expression of 0.42 - a 50% increase over 0.28 from ExPecto (**Figure S2)**.

Beyond the predictive performance of DEcode, we utilize the model to help define the biological processes regulating DE. Many studies have demonstrated that TFs-promoter interactions are critical determinants of transcriptional activity of promoter and thereby define gene expression levels^2^. However, it is unclear to what extent RNA features, which we define as each RNA’s binding sites of proteins and miRNA’s, contribute to gene expression levels compared to TFs-promoter interactions. To answer this question, we re-trained the deep learning model, randomizing either RNA features, promoter features, or both. We found that RNA features alone explained the actual TPM values better than the model trained with promoter features (**Figure 2E**). An example of how RNA features may distinguish between transcripts to a greater extent than promotor features can be seen in the structure of the gene ACADM (**Figure 2F**), which showed substantial differences between the promoter-based model and the RNA-based model. For instance, the promoter-based model could not distinguish 8 out of 11 transcripts coding for the ACADM gene that shared the same promoter region (p1 in **Figure 2F**). However, the actual expression levels for the 8 transcripts varied depending on the mRNA structures and therefore were more accurately captured by the RNA-based model (**Figure 2F**). However, the importance of RNA features was tissue-dependent (**Figure 2G**), as gene expression in the aorta and coronary arteries were mainly defined by TF-promoter interactions, whereas RNA-binding features were the major predictors for thyroid-specific or skeletal muscle-specific expression.

### Regulatory factors for differential expression across human tissues

To quantify the importance of the biological interactions weighted in the DEcode models, we calculated DeepLIFT scores, which are a measure of the additive contribution of its binding site to each prediction^14, 15^ and then averaged the DeepLIFT scores for each interactor across genes (**Table S2**). Because DeepLIFT scores for the gene-based model and the transcript-based model were well correlated (Spearman’s rho = 0.52, P < 2.2e-16) (**Figure S3**), we focused on DeepLIFT scores for the gene-based model in the following analyses. For the prediction of median TPM levels, the enrichment of the binding sites of RNABPs peaked among the top 12% of influential predictors, which was significantly greater than the influence of TFs and miRNAs (P < 0.00001) (**Figure 3A**). Indeed, out of the top 30 key predictors, 19 were RNABP’s binding sites and 11 were TF binding sites. The direction of DeepLIFT scores indicates either a positive or a negative effect of having the binding site on the abundance of RNA (**Figure 3B**). For instance, the binding sites of ATXN2, DDX3X, and FUS had high positive DeepLIFT scores to the prediction of RNA abundance, indicating the RNAs that bear binding sites for these RNABPs tended to be more highly expressed (**Figure 3B**). We also calculated DeepLIFT scores for tissue-specific expression to examine critical predictors for each of 53 tissues (**Figure 3C**). The DeepLIFT scores across tissues recapitulated the contribution of binding sites of known master regulators in each tissue such as REST for brain tissues^16^, SPI1 and RUNX1 for immune-related tissues^17^, TP63 and KLF4 for skin^18^, HNF4A for liver^19^, and PPARG for adipose-related tissues^20^, which suggested the differences in predictive contributions of binding sites of a given regulator reflect the differential activities of regulators across tissues. We hypothesized that the differential activities of a regulator could be in part explained by the relative abundance of a regulator across tissues. Based on this hypothesis, we contrasted DeepLIFT scores for the binding sites of each regulatory factor and its expression levels across tissues. We indeed found that 99 RNABPs and 410 TFs showed significant correlations between DeepLIFT scores of their binding sites and their expression levels (FDR < 5%) (**Figure S4**). These relationships were not based on differences in expression profiles between brain and non-brain tissues, as the relationships remained the same without brain tissues (**Figure S5**). The sign of the correlation possibly reflects whether the binding of a regulator to RNA increased or decreased the abundance of the RNA. For instance, the model suggested that PPARG and PTBP1 are positive regulators of gene expression as DeepLIFT scores of PPARG or PTBP1 binding sites were higher in the tissues expressing PPARG or PTBP1 at higher levels (**Figure 3D**). Indeed, PPARG is a transcriptional activator^20^ and PTBP1 is a stabilizer of RNAs^21^. Conversely, the expression levels of REST, a transcriptional repressor^16^, or METTL14, an RNA methyltransferase destabilizing RNAs^22^, showed inverse correlations with their DeepLIFT scores as expected (**Figure 3D**). These results indicated that DEcode reflects biological mechanisms for controlling RNA abundance.

**Figure 3.**
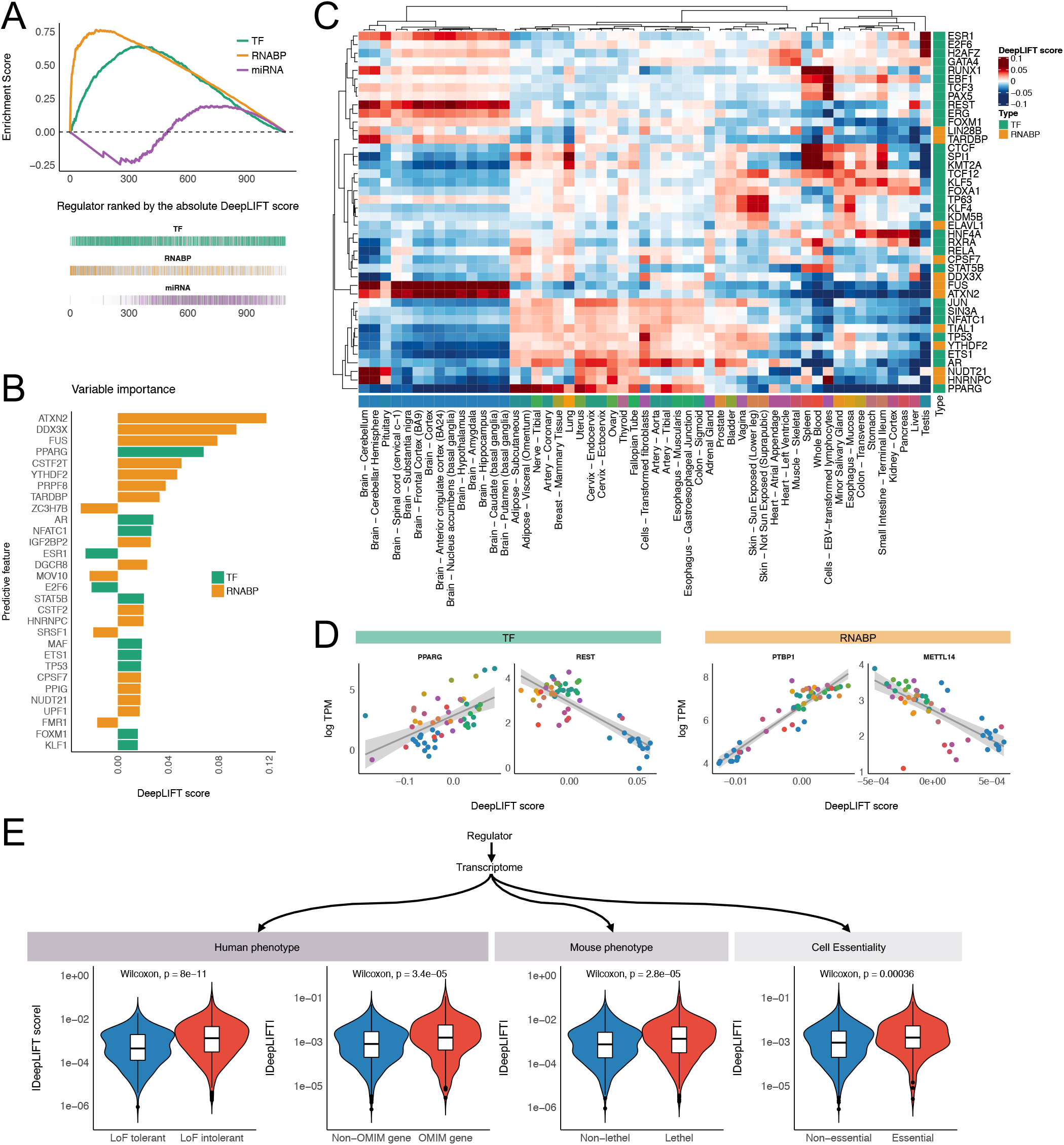
Identification and characterization of key predictors in the tissue-level models. (**A**) The enrichment of a regulator class in the key predictors for the median absolute expression levels. We ranked the regulators by the DeepLIFT scores and evaluated the enrichment of each regulator class. We used the pre-ranked gene set enrichment analysis (GSEA) algorithm with 10,000 permutations to compute enrichment scores and statistical significance. (**B**) Top 30 key predictors for the median absolute expression levels. (**C**) Key predictive regulators for the tissue-specific transcriptomes. We selected the top 5 key predictors for each tissue and their DeepLIFT scores were displayed as a heatmap. The ward linkage method with the Euclidean distance was used to cluster tissues and predictors. (**D**) Example relationships between the predictive importance for a regulator and its expression levels across tissues. (**E**) The overlap between the key regulators for the median absolute expression levels and external functional gene sets.

### Critical predictors of transcriptome are enriched for disease genes

Next, we characterized the roles of the critical regulators of human transcriptome, as suggested by the DEcode models (**Figure 3A**). We hypothesized that if these are truly impactful transcriptome regulators, then defects in such regulators would have significant impacts on cellular phenotypes and thereby lead to disease. To examine this hypothesis, first, we obtained genes whose loss-of-function (LoF) mutations are depleted through the process of natural selection, from the Exome Aggregation Consortium (ExAC)^23^. Since these genes are intolerant to LoF mutations they are considered to play important roles in individual fitness. Out of all TFs and RNABPs used in DEcode, 853 genes were examined in the ExAC study and 601 genes were reported as being intolerant of homozygous or heterozygous LoF mutations, with probability greater than 99%. We found that these LoF-mutation-intolerant regulators had greater DeepLIFT score magnitudes for the prediction of the absolute gene expression (**Figure 3E** and **Table S3**). In particular, these associations are based on genes that are intolerant to both heterozygous and homozygous LoF mutations (**Figure S6**). This suggested that having LoF mutations only in a single allele of the predicted critical regulators would cause a deleterious consequence on survival or reproduction in humans. Next, to examine whether the predicted critical regulators of transcriptome indeed cause diseases, we obtained disease-causing genes registered in the Online Mendelian Inheritance in Man (OMIM)^24^. We confirmed that mutations in the regulators with high DeepLIFT scores tended to cause genetic disorders (**Figure 3E**). Interestingly, their roles on fitness are likely preserved across species, as dysfunctions of the predicted critical regulators also led pre-weaning lethality in mice (**Figure 3E**). Lastly, we asked whether the loss-of-function of the predicted critical regulators of the transcriptome could also impair cellular viability, by overlapping them with loss-of-function screens for a range of cellular models, from the Cancer Dependency Map project (DepMap)^31^. We found that the key genes for cellular viability tended to have higher DeepLIFT scores in the DEcode model (**Figure 3E**). These results were robust, as they were also supported by the DeepLIFT scores for the transcript-level model (**Figure S7**). Together, the results indicated that the critical predictors of transcriptome indeed play critical roles in maintaining vital cellular and body functions. Thus the DEcode model can identify disease-causing genes, and this capability points toward the broader validity of predicted key regulators.

### DEcode predicts differential expression across individuals

Next, we asked whether the same input of promoter and RNA features could also predict relative expression differences across individuals within the same tissue. We hypothesized that each individual has different activation levels of regulatory factors, and thus those differences lead to person-specific differential expression of their targets. To verify our hypothesis, we extended the DEcode framework to model differential expression across individuals for 14 representative tissues with a sample size greater than 100 in GTEX. This was challenging as the average variance in gene expression within tissues was less than 25% of that between tissues (**Figure S8**).

To generate person-specific predictions, we utilized transfer learning, wherein the parameters in convolutional layers in the across-tissue DEcode model were fixed and then only the parameters in the fully-connected layers were tuned (**Figure S9**). The person-specific models successfully predicted fold changes across individuals with a mean Spearman’s correlation of ~0.28 (**Figure 4A**). The performance was further increased to 0.34 when we filtered out the models that worked poorly for the validation data (**Figure S10**). Note the model selection was performed based on validation data alone, and all the follow-up performance evaluations and analyses were conducted by using testing data to prevent information leaks that could inflate model performance (**Figure S9)**. The models were indeed person-specific as they did not predict gene expression profiles of unrelated individuals (**Figure 4A**). To examine if the model captured the person-specific expression shared across tissues^25^, we compared expression between tissues within the same individuals and between different individuals. The predicted expression showed better concordance between tissues from the same individuals, as is the case with actual expression data, which indicated the model captured the person-specific regulatory mechanisms, even though we did not use any direct information that could identify individuals (**Figure 4B**).

**Figure 4.**
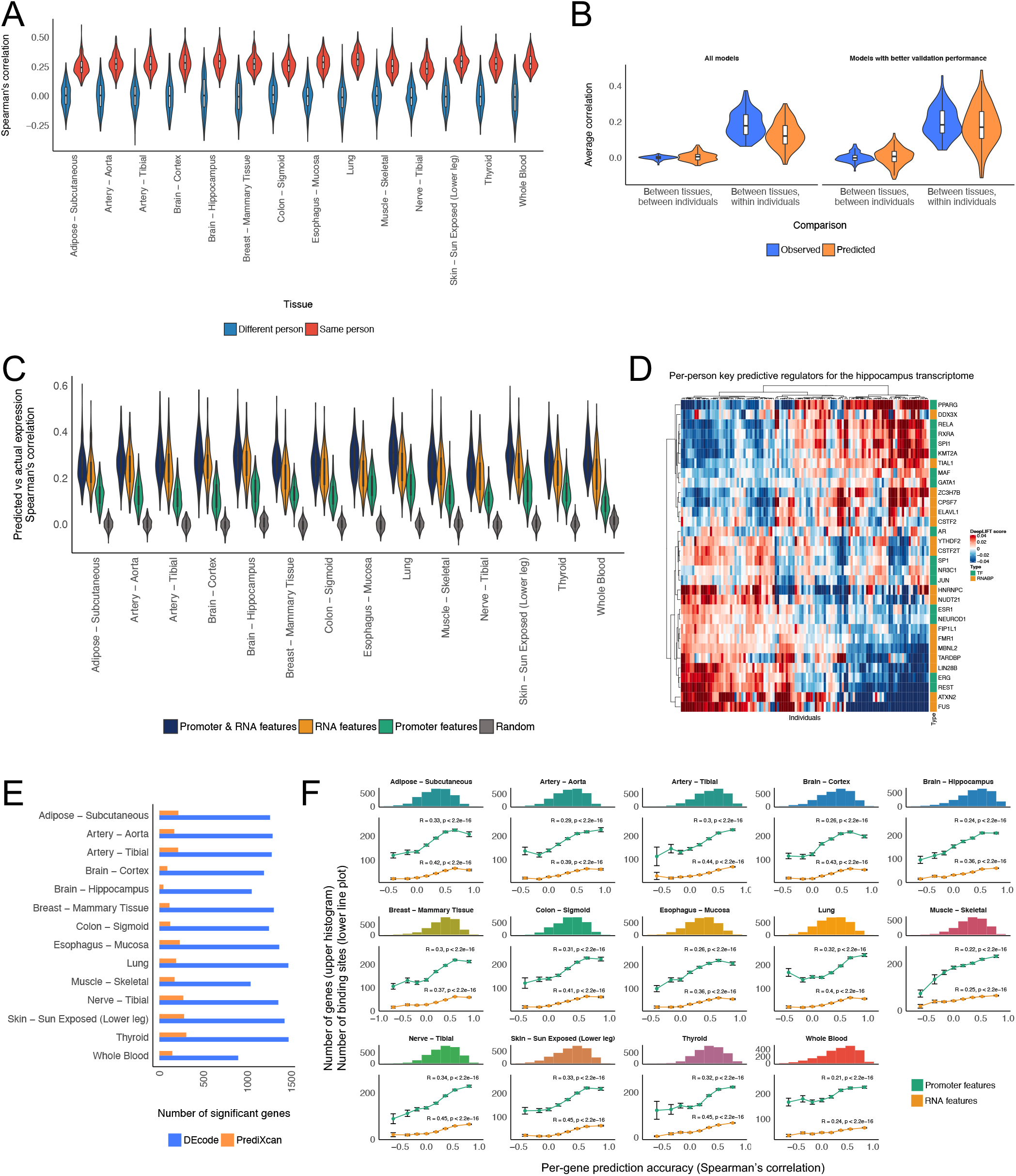
Performance of the person-specific models. (**A**) The predictive performances of the person-specific models for the actual data from the same individuals and unrelated random individuals. (**B**) The person-specific models predicted person-specific expression shared across tissues. (**C**) The performances of the models trained with a distinct feature set. (**D**) Per-person key predictive regulators for the hippocampus transcriptome. We selected the top 5 key predictors of the hippocampus transcriptome for each individual and their DeepLIFT scores were displayed as a heatmap. The ward linkage method with the Euclidean distance was used to cluster tissues and predictors. (**E**) Comparison of per-gene predictive accuracy between DEcode and PrediXcan. The number of genes that showed a positive Pearson’s correlation between predicted and actual gene expression levels at FDR 5% was calculated for each method. Only the testing genes on chromosome 1 were used for this comparison. (**F**) Per-gene prediction accuracy is associated with the number of features present in RNAs and promoters. The histogram represents Pearson’s correlations between the predicted and the actual expression for each gene. The line plot shows the average number of RNA and promoter features of genes in each bin of the histogram. Spearman’s correlation between the number of features and per-gene correlations is displayed in the line plot. The error bars indicate standard errors.

Next, to gauge the contribution of RNA and promoter features to the person-specific expression profiles, we re-trained models with randomized RNA features, promoter features, or both. The RNA-feature-based model performed on average 85% as well as the model trained with all features. This corresponded to an average 173% performance gain, compared to the promoter-feature-based model, which suggested that the post-transcriptional controls are the major determinants of the differential expression across individuals (**Figure 4C**). The model also allowed us to investigate the person-specific activities of regulators by calculating DeepLIFT scores (**Figure 4D**). At least 100 of regulators out of 933 regulators in each tissue showed a good correlation between their DeepLIFT scores and expression levels across individuals (**Figure S11**). The signs of these correlations were consistent between tissues, and consistent with those of the cross-tissue model (**Figure S12**). This suggested that differential expression between individuals and between tissues can be modeled by the universal relationships between regulators and their targets.

To examine whether specific genes contributed to the per-person accuracy of the predicted gene expression, we also assessed its accuracy on a per-gene basis. The predicted expression of a majority of the testing genes (78% on average) showed significant positive correlations with the actual gene expression (FDR<5%). In order to assess whether this predictive performance outperformed a state of the art method, we compared DEcode with PrediXcan^26^, which predicts person-specific gene expression from genetic variations in cis-regulatory regions of genes. We built PrediXcan models for each of the testing genes based on the same GTEX gene expression data used for the DEcode models and whole-genome sequence data of corresponding individuals (see Methods). The PrediXcan model predicted gene expression levels of only about 11% of the testing genes at FDR less than 5%, which was far less than that of DEcode (**Figure 4E**). This suggested that the differential activity of transcriptional and post-transcriptional regulators has a larger effect on gene expression than genetic variations in cis-regulatory regions.

The genes that DEcode could predict well were similar across tissues (**Figure S13**). This suggested that the predictability of gene expression is defined by gene characteristics rather than a target tissue. We, therefore, explored gene characteristics that were associated with the per-gene accuracy of the predicted expression. We found that the models showed higher performance for the genes that are registered in multiple gene annotation databases than those found only in the GENCODE database (**Figure S14**). The GENCODE-specific genes are novel or putative and thus their annotations are not well established. Since both actual gene expression and binding features in RNA and promoter regions are likely to be less accurate for such a novel or putative gene, it is reasonable that the performance of the model for those genes was lower than other well-established genes. Beyond the annotation reliability, we found that the number of known binding features for each gene had a larger effect on the predictability (**Figure S15**). This suggested that the more information on RNA and promoter interactions is available, the more the prediction becomes accurate. Interestingly, the number of binding features in RNAs was a stronger determinant of the predictive accuracy than that in promoter regions (**Figure 4F** and **Figure S15**). RNA-protein interactions are largely missing as global RNA-binding profiles are available for only about 10% of known RNABPs^30^. Thus, the incompleteness of RNA features is likely to be an origin of lower accuracy for a portion of genes.

### DEcode predicts trait-related transcriptomic changes

Next, we asked whether the person-specific expression profiles predicted by the DEcode models also retained trait-associated differential expression changes. For this, we conducted differential expression analysis against the donor’s age and sex using the predicted gene expression data. Notably, test statistics of the predicted data showed significant positive correlations with those of the actual data in all tissues for both traits (**Figure 5A**). Especially, age- and sex-specific expression changes were well preserved in the predicted data in lung (Spearman’s rho = 0.59, P < 2.2e-16) and hippocampus (Spearman’s rho = 0.47, P < 2.2e-16), respectively. The predicted associations were the closest to those of corresponding tissues in 9 and 11 out of 14 tissues for age and sex, respectively (**Figure 5B**). This indicated that the predicted gene expression changes against age and sex are tissue-specific in most cases, rather than the effects shared across tissues. We also explored the regulators for the age- and sex-related gene expression changes by associating regulator’s DeepLIFT scores with age and sex. We found that many regulators, for instance, 717 in the tibial artery and 904 in the breast mammary tissue, showed age- and sex-dependent changes at FDR 5%, respectively (**Figure 5C and Table S4**), which showed the capability of DEcode to associate transcriptional regulators with phenotypes. Although there were more TFs associated with phenotypes than RNABPs and miRNAs, overall collective impacts of RNA features on the generative process of DEs for age and sex were greater than those of promoter features in most tissues (**Figure S16)**.

**Figure 5.**
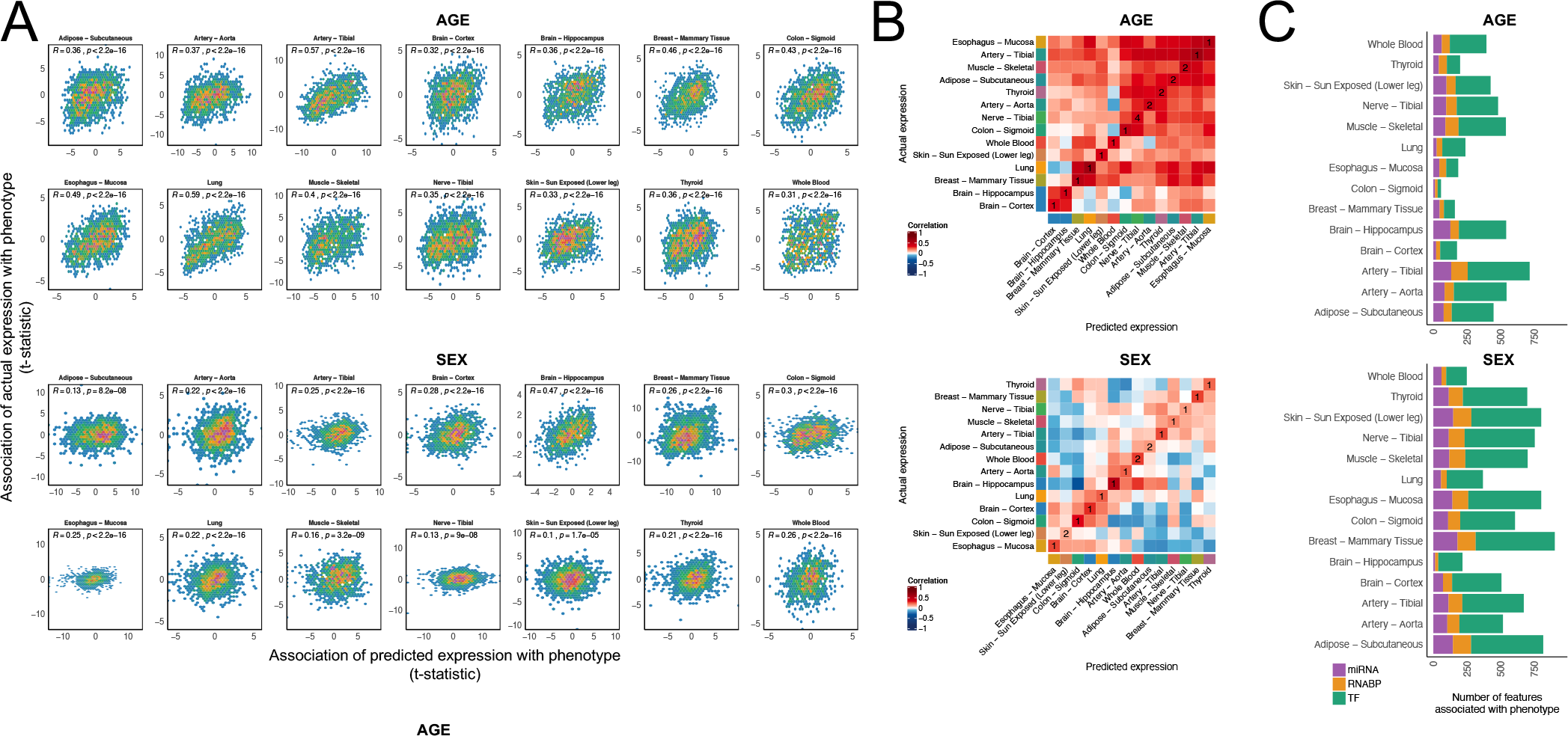
Application of the person-specific models to analyze phenotype-related gene signatures. (**A**) The scatter plots showing the relations between the associations of genes with age and sex using predicted expression and those using the actual expression. Spearman’s correlation between t-statistics using the predicted and the actual gene expression is displayed in the scatter plot. (**B**) The pairwise Spearman’s correlations between the predicted and the actual associations of genes with age and sex in all tissues. The numbers in diagonal elements of the heatmap indicate the ranks of similarity of the predictions with the actual observations in the corresponding tissues. (**C**) The regulators whose DeepLIFT scores were associated with age and sex. The Benjamini–Hochberg procedure was used to control the false discovery rate at 5% for each phenotype.

### DEcode predicts gene co-expression relationships

Co-expression analysis is a frequent component of transcriptome studies as gene-to-gene co-expression relationships are regarded as functional units of the transcriptional system^27^. Therefore, we examined if the DEcode models could detect known gene co-expression relationships. These tests were both a potential validation of the person-specific DEcode predictions, and a means to explore the biological basis of co-expression. We found that the gene co-expression relationships in the predicted gene expression profiles separated gene pairs with positive and negative correlation in the actual gene expression data in each tissue (**Figure 6A**). Furthermore, the predicted gene expression profiles also detected inter-tissue co-expression relationships (**Figure 6B**). The accuracy of these results motivated us to investigate key factors driving co-expression, via the DEcode predictions. RNA features alone could explain co-expression relationships better than promoter features in most tissues (**Figure 6C**), which again suggested the significant contribution of RNA features to person-specific transcriptomes.

**Figure 6.**
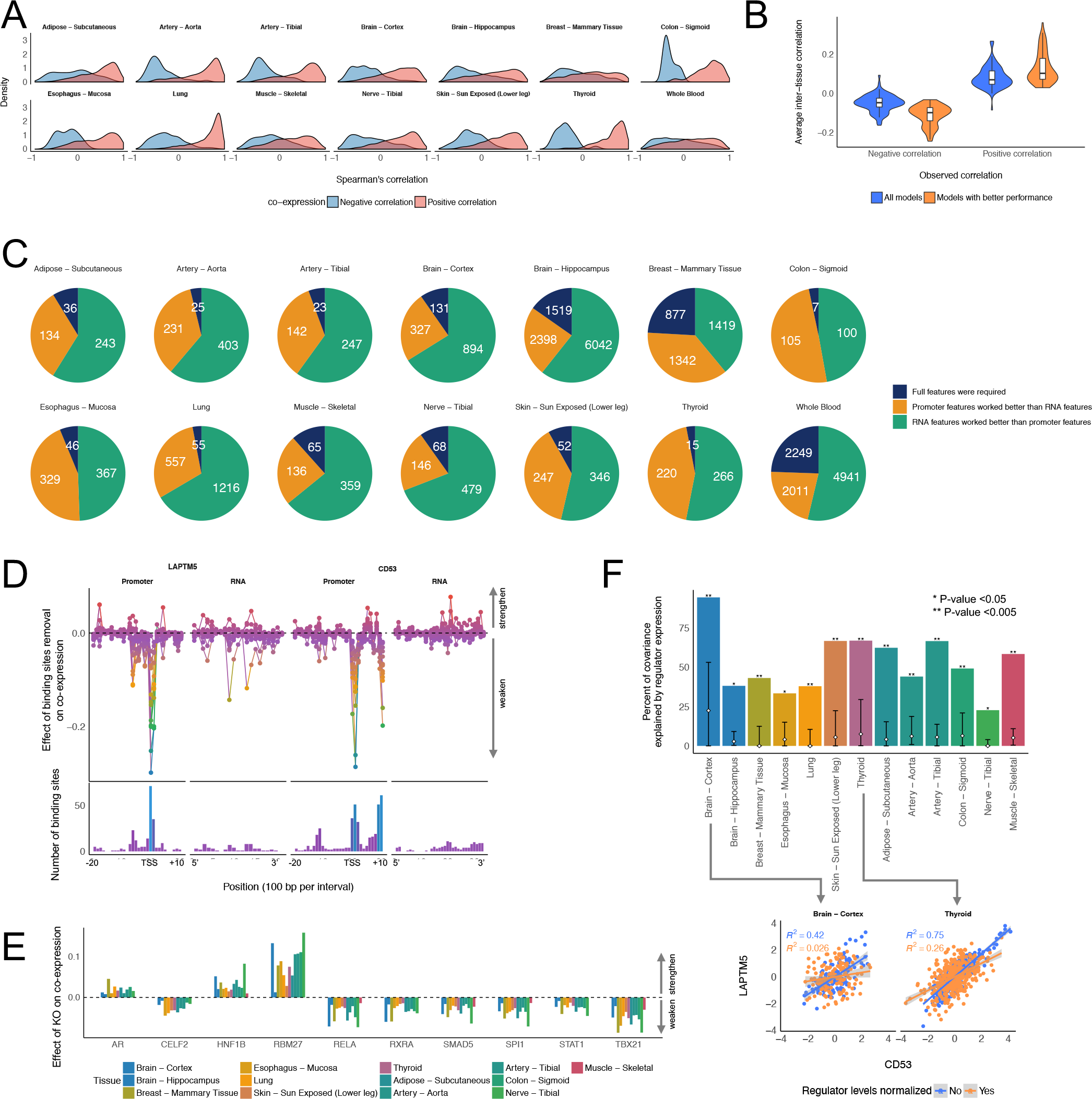
Application of the person-specific models to investigate gene co-expression. (**A**) Co-expression relationships in the predicted gene expression. We defined the ground truth co-expression relationships as gene pairs with the absolute Spearman’s correlation greater than 0.7 in the actual expression of the testing data. The density of Spearman’s correlation between the co-expressed gene pairs in the predicted gene expression data was estimated using the density function in R with the Gaussian kernel. (**B**) Inter-tissue gene co-expression relationships in the predicted gene expression. We defined the ground truth co-expression relationships as gene pairs with the absolute Spearman’s correlation greater than 0.5 in the actual expression of the testing data. We computed the average of Spearman’s correlation of the inter-tissue co-expressed gene pairs in each pair of tissues using the predicted expression from all models and the models whose performances on validation data were greater than 50% percentile in each tissue. (**C**) The major feature types contributed to the gene co-expression. We defined the gene pairs with the absolute Spearman’s correlation greater than 0.3 and the sign of the correlation matched with one with the ground truth as the successfully predicted gene pairs. The successfully predicted gene pairs of the model trained with the full set of features were split into three groups based on the performance of the models trained with only RNA features or promoter features. (**D**) The effect of the binding site removal on co-expression between *LAPTM5* and *CD53*. We simulated gene expression profiles with random removals of the binding sites in each gene 10,000 times and computed a correlation between *LAPTM5* and *CD53* for each simulation. Multiple regression was used to estimate the effect of the binding site removal on the co-expression in each tissue. (**E**) The key regulators for the co-expression between *LAPTM5* and *CD53*. We simulated gene expression profiles with random KOs of regulators in each gene 10,000 times and computed a correlation between *LAPTM5* and *CD53* for each simulation. We used multiple regression to estimate the effect of the KO on the co-expression in each tissue and identified the consensus regulators across tissues. (**F**) Percent of the co-expression relationship explained by the expression levels of the key regulators. The white diamond and the error bars in the bar indicated the average and 95% percentile of the percent of variance explained by randomly picked regulators, respectively. The scatter plots show the effect of the key regulators on the co-expression.

To further assess the capability of DEcode to decipher the mechanisms leading a specific co-expression relationship, we focused on the co-expression of *LAPTM5* and *CD53*, which were robustly co-expressed both in the simulated expression data and the actual data in all tissues except whole-blood. Using the trained model, we simulated the consequences of disruptions of promoter and mRNA features. The co-expression relationship was weakened when the features near transcriptional start site (TSS) and 1,000 bp downstream of TSS in *LAPTM5* or near TSS and 500 bp upstream of TSS in *CD53* were removed (**Figure 6D**). These observed effects were reasonable because many TFs bind to these regions (**Figure 6D**). We further examined the specific regulators for the co-expression relationships by simulating knockout (KO) effects of regulators. The *in-silico* KO experiments revealed that immune-related TFs such as *SPI1* and *TBX21* potentiated the co-expression relationships consistently across multiple tissues (**Figure 6E**). To validate if these regulators indeed induced the co-expression relationships, we conducted a mediation analysis that is an orthogonal computational method to infer the effect of regulators on downstream targets. A mediation analysis evaluated the hypothesis where if *LAPTM5* and *CD53* are co-expressed due to the predicted regulators, normalizing expression levels of the two genes by the expression levels of the regulators would decrease the co-expression relationships. Specifically, it quantified the covariance between *LAPTM5* and *CD53* explained by the expression levels of the predicted regulators using the actual expression data. The set of the 10 regulators together mediated up to 94% of covariance, which was significantly greater than the same number of randomly picked regulators (**Figure 6F**). This example showed the utility of DEcode framework to identify the drivers of the co-expression.

### DEcode reveals molecular regulations for frequently DE genes in meta-transcriptomes

A recent meta-analysis of over 600 human transcriptome data revealed that some genes are more likely to be detected as DE genes than others in diverse case-control studies^28^. From this observation, Megan et al. formulated the “DE prior”, a global ranking of gene’s generic likelihood of being DE. The genes with high DE prior rank were significantly more enriched with DE genes from a variety of conditions, as compared to other functional gene sets, such as those contained in gene ontology or canonical pathways^28^. However, the regulatory-origin behind the ranking of these highly responsive genes has yet to be uncovered. Therefore, we used DEcode to examine whether the DE prior rank could be generated by gene regulatory interactions, and to identify critical regulatory relationships for frequently DE genes. The ability of DEcode to predict global DE prior ranks was highly significant (P < 2.2e-16) and practically relevant (Spearman’s rho = 0.53) (**Figure 7A**). Furthermore, DEcode was able to identify genes with high (90th percentile and greater) DE prior probability (AUCROC = 0.81, 95% confidence interval = 0.78 - 0.84) (**Figure 7B**). Re-training the model with randomized inputs indicated that TF-promoter interactions were the major factors explaining the DE prior rank (**Figure 7B**). To further characterize TFs that contributed to the prediction, we defined TFs with DeepLIFT score greater than 90th percentile as critical TFs (**Table S5**) and performed pathway analysis on them. We found that critical TFs were enriched for cancer or inflammatory-related KEGG pathways (FDR<5%) such as pathways in cancer (Fold = 3.1, P = 4.2e-5), JAK-STAT signaling pathway (Fold = 6.8, P = 4.8e-5), chemokine signaling pathway (Fold = 7.3, P = 1.4e-4), and acute myeloid leukemia (Fold = 4.5, P = 3.6e-4) (**Table S6**). This result is consistent with the disease-related data context for DE prior, which is 62% cancer-related and 23% inflammatory-related. Supported by the ability to predict DE prior ranks, and by the consistency of these results, this application of DEcode illustrates how it goes beyond DE gene lists, to uncover major key drivers for generating DE.

**Figure 7.**
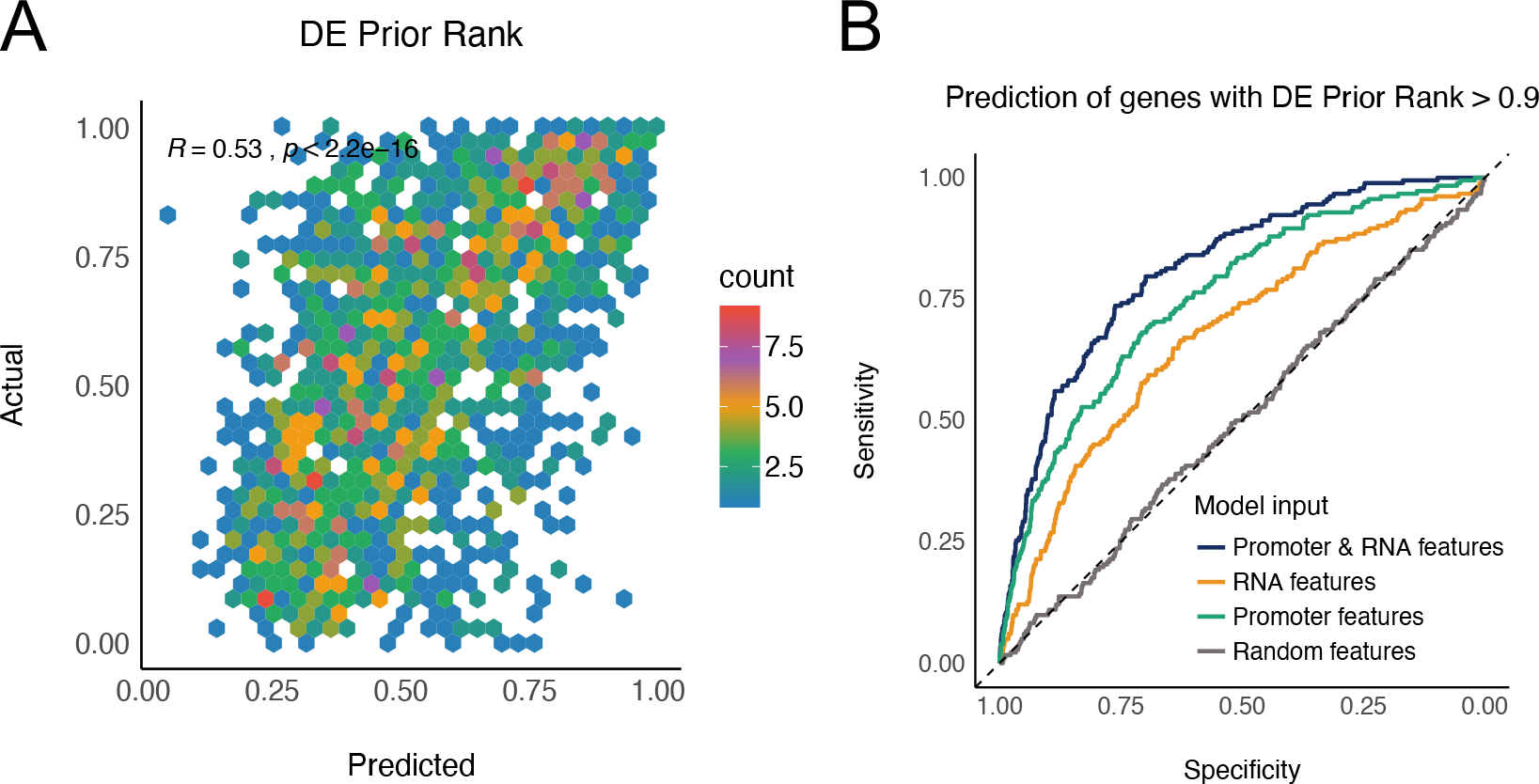
DEcode predicts gene’s prior probability of differential expression. (**A**) The scatter plots showing the relations between predicted and actual DE prior rank. The predicted logit of DE prior rank was converted to probability and compared with actual DE prior rank with Spearman’s correlation. (**B**) The performances of the models trained with a distinct feature set. ROC represents the performance of model predicting genes with DE prior rank greater than 0.9.

In summary, DEcode defines major principles in gene regulation in arbitrary gene expression data. It is applicable to tracing the origins of complex gene expression patterns such as co-regulation, and also to arbitrary gene expression signatures. This capacity is strongly supported on a comparative basis to alternative methods, and on an absolute basis across diverse applications, which include, through predictions of transcript-usage, person-specific gene expression, frequently DE genes of multiple external disease-related gene sets.

## Discussion

We introduced the DEcode framework, which integrates a wealth of genomic data into a unified computational model of transcriptome regulations to predict multiple transcriptional effects, including the absolute expression differences across genes and transcripts, tissue- and person-specific transcriptomes. Systems biology analysis of these results provided biological insights regarding the regulatory mechanisms of transcriptome. For instance, it suggested that absolute expression levels are mainly under post-transcriptional control, whereas tissue-specific expression is shaped by both transcriptional and post-transcriptional control. This implied that TFs act as a switch that initiates tissue-specific transcriptional programs, but once a gene is transcribed at a certain level, its abundance in the cells will be primarily regulated by RNABPs. The post-transcriptional regulators were also critical for explaining individual differences in transcriptomes and thus may fine-tune the transcriptome in response to environmental and genetic factors.

Transcriptome analysis often identifies differentially expressed genes and then assesses the enrichment of functional genes such as TF-targets one by one. The person-specific DEcode model offers several comparative advantages. First, DEcode can take into account the effects of multiple regulators simultaneously as opposed to one at a time. Second, DEcode can estimate the person-specific regulator’s activities that can be used to identify regulators associated with a phenotype of interest. Third, DEcode can simulate the consequence of KO perturbations for each gene. This step can reduce the number of candidate key drivers of gene expression changes by an order of magnitude or more, and facilitates the design of follow-up experiments. Therefore, DEcode can extract more actionable information from transcriptome data, which will benefit a variety of transcriptome studies.

Looking toward even more expansive applications, the DEcode framework has the flexibility to incorporate other types of genomic information such as DNA methylation, histone marks, and RNA modifications, and also can be extended to other organisms. Thus, DEcode framework provides a direct bridge between accumulating genomic big data and individual transcriptome studies, allowing researchers to predict molecules that control DE associated with any condition or disease.

## Materials and Methods

### Transcriptome data processing

To prepare gene expression data used for the model training, we downloaded the median gene TPM from 53 human tissues from the v7 release of GTEX portal (https://gtexportal.org). We kept 27,428 genes expressed greater than two TPM in at least one tissue and log2-transformed TPM with the addition of 0.25 to avoid a negative infinity. Then, we calculated the median log2-TPM across 53 tissues and log2-fold-changes relative to the median of all tissues. The processed gene-level expression data comprised 27,428 genes with 54 columns including relative fold-changes for 53 tissues and the median log2-TPM across 53 tissues. To compile transcript-level data, we downloaded the individual-level transcript TPM from the GTEX portal and computed the median transcript TPM by tissue. We processed the transcript data in the same way we did for the gene-level data. The resulted transcript-level data included 79,647 transcripts that corresponded to 23,813 genes. For building person-specific DEcode models, we obtained the gene-level TPM for each individual in 14 tissues from the GTEX portal. We filtered out lowly-expressed genes in each tissue and kept genes expressed greater than one TPM in at least 50% of samples. Then, we log2-transformed TPM with the addition of 0.25 and then quantile normalized the log2-TPM. Finally, we removed the effects of technical covariates including rRNA rate, intronic rate, and RIN number via linear regression for each gene followed by quantile normalization.

### Promoter and RNA binding features

To generate RNA and DNA feature matrices, we downloaded genomic locations of binding sites of 171 RNABPs from POSTAR2^29^ as of Oct 2018, 218 miRNAs from TargetScan Release 7.2^30^, and 826 TFs from GTRD^31^ as of Oct 2018. Then, we mapped the binding sites of RNABPs, miRNAs, and TFs to promoters and RNA-coding regions defined in the GTF file provided by the GTEX portal. A promoter region of each gene was defined as the region from 2,000 bp upstream of the transcriptional start site (TSS) to 1,000bp downstream of the TSS. We only used interactors that bind to promoters or RNA-coding regions of at least 30 genes, or transcripts as the predictors in each model. To reduce the size of the input, an RNA-coding region and a promoter region of each gene was binned with 100 bp intervals and the number of bases bound to each RNABP, miRNA, or TF was counted in each interval. This step generated RNA and DNA feature matrices for each gene described in **Figure 1**.

### Training tissue-specific models

For training the gene-level model of tissue-specific expression, we reserved all 2,705 genes coded on chromosome 1 as the testing data and the rest of the genes was randomly split into training data (22,251 genes) and validation data (2,472 genes). In the case of the transcript model, we used all 7,631 transcripts coded on chromosome 1 as the testing data and the rest of the transcripts was randomly split into training data (64,978 transcripts) and validation data (7,038 transcripts). The binding matrices were normalized by the maximum values for each binding protein and miRNA. The relative fold-changes for 53 tissues were scaled together to set the standard deviation as one and the median log2-TPM was separately scaled to set the standard deviation as one. These steps were conducted for the training data first and then the same scaling factors were used for the validation and the testing data to avoid information leaking from those data. We constructed and trained DL models using Keras (version 2.1.3)^32^ with a TensorFlow (version 1.4.1)^33^ backend. Hyper-parameters were optimized using hyperopt (version 0.2)^34^ based on the mean squared error against the validation data. The detailed structure of the model was described in **Figure S17**. The training was done using mini-batches of 128 training examples with a learning rate of 0.001 for Adam optimizer^35^. The number of maximum training epochs was set to 100 with early-stopping of 10 based on validation loss. This training cycle was repeated 10 times and the best model for the validation data was selected as the final model (**Figure S1**). All models were trained using TITAN X Pascal graphics processing units (Nvidia).

### Comparison of DEcode with ExPecto

To perform a fair comparison between DEcode and ExPecto^11^, we used 18,550 genes that were commonly included in both studies and trained models with the same set of genes for training and evaluation. Since ExPecto model was originally built using genes on chromosome 8 as the testing data, we followed the same procedure as we reserved all 714 genes coded on chromosome 8 as the testing data and the rest of the genes was randomly split into training data (16,052 genes) and validation data (1784 genes). The epigenetic states estimated by ExPecto were downloaded from the ExPecto repository (https://github.com/FunctionLab/ExPecto) as of Nov 2019. Given the epigenetic states, we built a prediction model for tissue-specific gene expression for each tissue via XGBoost based on the training script downloaded from the ExPecto repository. We modified the original script so that the early stopping of the model optimization was decided based on the performance on the validation data instead of the testing data. This modification prevented the overfitting of the model to the testing data. We used the same hyper-parameters for XGBoost as in the script. Both hyper-parameters and model parameters of DEcode model were also trained with the same set of training, validation, and testing genes.

### DeepLIFT score calculation

To evaluate the importance of input features to the prediction, we calculated DeepLIFT (Deep SHAP) scores^14^ using DeepExplainer implementation (version 0.27.0)^15^. The DeepLIFT method estimates the contribution of each input compared to a reference input in a trained DL model. To compute the contribution of the presence of a binding site, we used a reference that does not have any binding sites in both promoters and RNAs with the median length of all genes in the testing data. DeepLIFT scores follow a summation-to-delta property where the summation of input contributions (DeepLIFT scores) is equal to the difference in the predicted value compared to the prediction from the reference input. We calculated DeepLIFT scores for each gene in testing data for each of 54 outputs, then summed up the scores over promoter or RNA regions for each feature, and finally averaged them over genes.

### Disease genes

The probability that a gene is intolerant for a loss-of-function mutation was downloaded from the release 1.0 of the ExAC portal (http://exac.broadinstitute.org). Disease genes were obtained from the OMIM portal as of June 2019 (https://www.omim.org/). We excluded provisional gene-to-phenotype associations and genes associated with non-disease phenotypes, multifactorial disorders, or infection. We obtained mouse-lethal genes from Gene Discovery Informatics Toolkit (v1.0.0)^36^ that provided pre-processed gene lists from the murine knock-out experiments registered in Mouse genome informatics (MGI)^37^ and the International Mouse Phenotyping Consortium (IMPC)^38^. The results of CRISPR screening for the genes essential for proliferation or viability conducted in the DepMap project^39^ were downloaded from Enrichr portal^40, 41^ as of June 2019 (https://amp.pharm.mssm.edu/Enrichr). Enrichr portal provided two CRISPR screening results conducted independently at Broad Institute and the Sanger Institute. To reduce the false positives in the CRISPR screening, we used essential genes that were identified in both of the two independent screenings.

### Training person-specific models

To train person-specific models, we utilized the same model structure as the tissue-model, except that the number of model outputs was modified to match the sample size of the tissue. We re-used the parameters of convolutional layers in the tissue-model and only parameters in the fully-connected layers were tuned (**Figure S9**). We used the same gene splits and the same procedure of normalization and scaling as the tissue-model for training and evaluating models. We evaluated the model prediction for each individual separately based on validation data and filtered out the individual models that performed less than 50% percentile of all individual models for some analyses (**Figure S9**).

### Training PrediXcan models

To build a prediction model for gene expression from genotype data, we trained PrediXcan^26^ models with GTEX gene expression and genotype data. A QCed vcf file of GTEx genotype data called by whole-genome sequence was downloaded from dbGaP for 635 individuals. We filtered out variants with a missing rate greater than 1% and minor allele frequency less than 1% and kept 9,219,660 variants for PrediXcan. We followed the model building procedure employed in PredictDB (http://predictdb.org/), a repository of PrediXcan models, as of Nov 2019. Briefly, we randomly split the samples into 5 folds. Then for each fold, we removed the fold from the data and used the remaining data to train an elastic-net model using 10-fold cross-validation to tune the lambda parameter. With the trained model, we predicted gene expression values for the hold out samples. We applied the PrediXcan method to predict the same gene expression data used for the person-specific DEcode models. We built PrediXcan model for each gene using variants located within 1 Mbp upstream and downstream of its TSS. A missing value of the genotype data was replaced with an average dosage of non-missing samples.

### Differential expression analysis for age and sex

Limma^42^ was used to identify genes associated with age using gender as a covariate. The log2-TPM values of genes in the testing data were used. We also tested the associations between DeepLIFT scores for predictors and age via limma to identify regulators for DE against ages and sex. The Benjamini–Hochberg procedure was used to control the false discovery rate at 5%.

### *in silico* binding-site disruption experiment

To simulate the consequence of the removal of binding sites on the expression of *LAPTM5* and *CD53*, we generated 10,000 synthetic inputs for each of *LAPTM5* and *CD53* where all binding sites in each interval of its promoter and RNA were randomly removed. From each of these synthetic inputs, we computed predicted expression values and correlated them with ones of another gene without any disruptions in its binding sites. Then, we used multiple linear regression to associate the location of disrupted regions with the correlation values between *LAPTM5* and *CD53* to estimate the effects of the disruption in each region on the co-expression relationship.

### *in silico* knockout experiment

To simulate the effect of regulator knockout (KO) on the expression of *LAPTM5* and *CD53*, we generated 10,000 synthetic inputs for each *LAPTM5* and *CD53* where each protein or miRNA bound to its promoter or RNA was randomly removed from it feature matrices. From each of these synthetic inputs, we computed predicted expression values and correlated them with ones of another gene without any removals in its feature matrices. Then, we used multiple linear regression to associate KOs of regulators with the correlation values between *LAPTM5* and *CD53* to estimate the effects of the KO of each regulator on the co-expression relationship. We applied the Bonferroni correction to control multiple testing and the regulators with the corrected p-value less than 0.05 in all tissues were chosen as the key drivers of the co-expression.

### Conditional independence test

To validate the effect of the predicted drivers on co-expression, we conducted a conditional independence test. We regressed the actual log2-TPM values of *LAPTM5* and *CD53* with the actual log2-TPM values of the predicted drivers and computed R^2^ (variance explained) between the residuals of two genes. The R^2^ based on the actual gene expression and one from the residuals were compared to quantify the covariance explained by the predicted drivers. To evaluate the significance of this effect, we repeated this process 1,000 times with an equal number of randomly picked genes that have a binding site in *LAPTM5* or *CD53* as regressors.

### DEcode model for DE prior rank

DE prior rank was downloaded from https://github.com/maggiecrow/DEprior. In the DE prior rank, each gene has a probability-like value where zero is the minimum and one is the maximum. To convert this value to a non-bounded scale, we applied the logit transformation to the DE prior value. We assigned a value of 10 to a gene that had an infinite value after the logit transformation. We used the same gene splits as the GTEX-tissue-model, which resulted in 13,433 genes for training, 1,504 genes for validation, and 1,674 genes for testing. We trained the DEcode model for DE prior rank using the same procedure as with the GTEX-person-specific models. To evaluate the contribution of promoter and RNA features to the prediction, the model was also trained with randomized input features. Receiver operating characteristic (ROC) curve analysis was performed using pROC R package^43^ with a default setting. We performed pathway analysis of the TFs with a DeepLIFT score greater than 90th percentile using KEGG pathways^44^. KEGG pathway gene sets were downloaded from MSigDB v6.1^45^. The enrichment significance was based on results of the hypergeometric test, with 757 unique TF genes as a background, against KEGG pathways comprised of at least 5 background genes. FDR was controlled at 5%. We manually curated the 159 disease-related data sets used in the construction of the DE prior ranking, to determine the number of data sets related to cancer or inflammatory disease.

### Code and model availability

DEcode software and pre-trained models for tissue- and person-specific transcriptomes are available at www.differentialexpression.org.

## Acknowledgments

We thank Dr. Lei Yu for managing access to GTEX data. The study was supported by NIH grants P30AG010161, R01AG061798, and R01AG057911. The Genotype-Tissue Expression (GTEx) Project was supported by the Common Fund of the Office of the Director of the National Institutes of Health, and by NCI, NHGRI, NHLBI, NIDA, NIMH, and NINDS. The data used for the analyses described in this manuscript were obtained from the GTEx Portal on 10/01/2018.

## Author Contributions

ST contributed to the conception and design of the study. ST performed the computational analysis. ST, CG, SM, and YW interpreted the result. ST wrote the first draft of the manuscript. All authors contributed to manuscript revision, read and approved the submitted version.

