## Supplementary figures and images for "Decoding differential gene expression"

### Figure S1

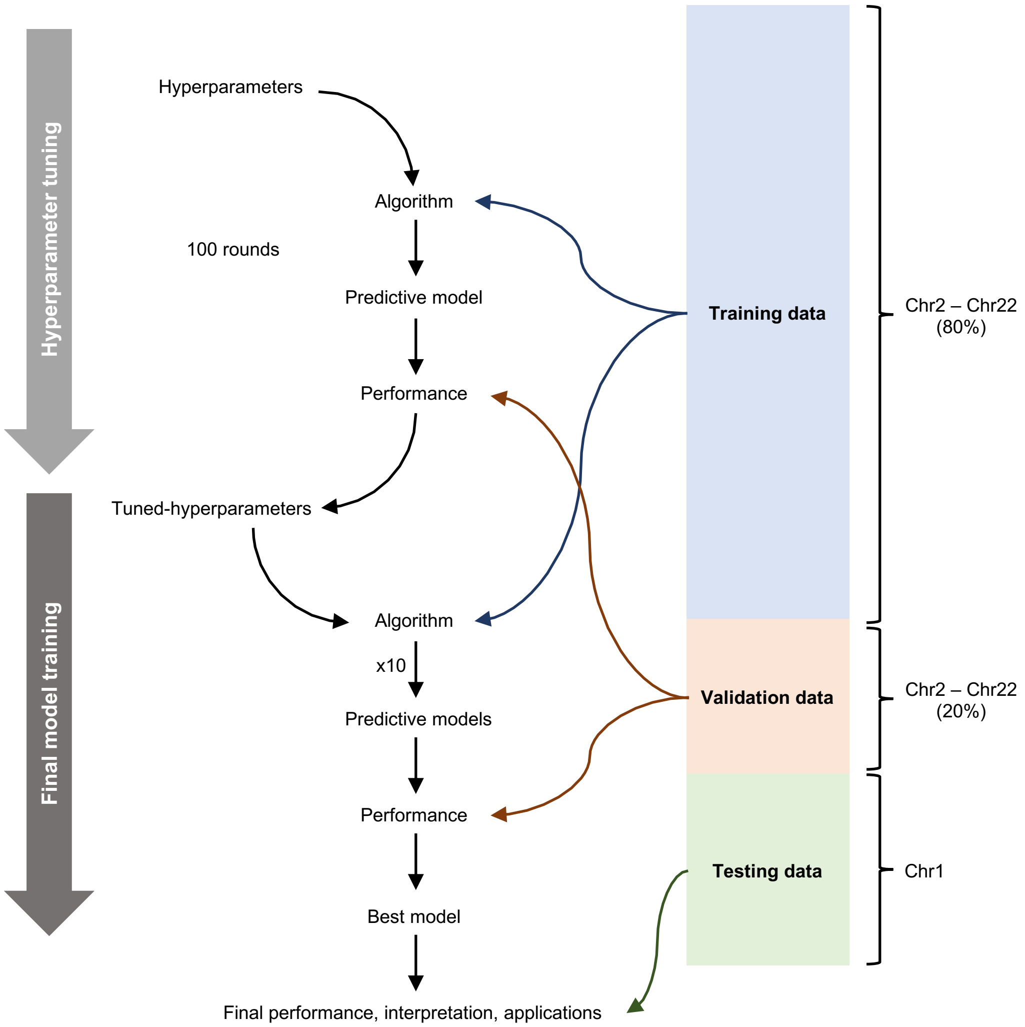


**Figure S1**. Overview of DEcode model building pipeline for tissue-specific expression.

### Figure S9

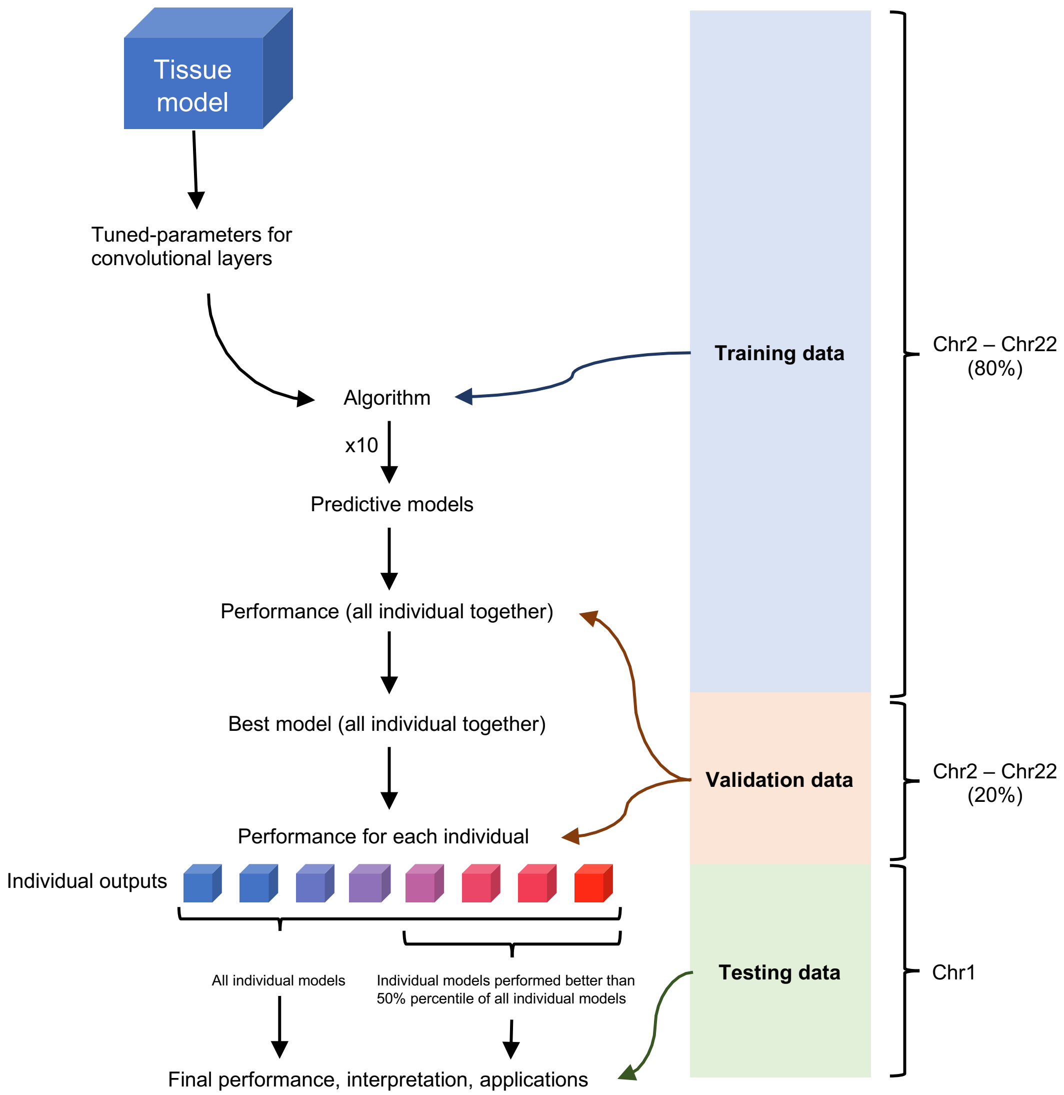


**Figure S9**. Overview of DEcode model building pipeline for person-specific expression.

### Figure S17

Gene model

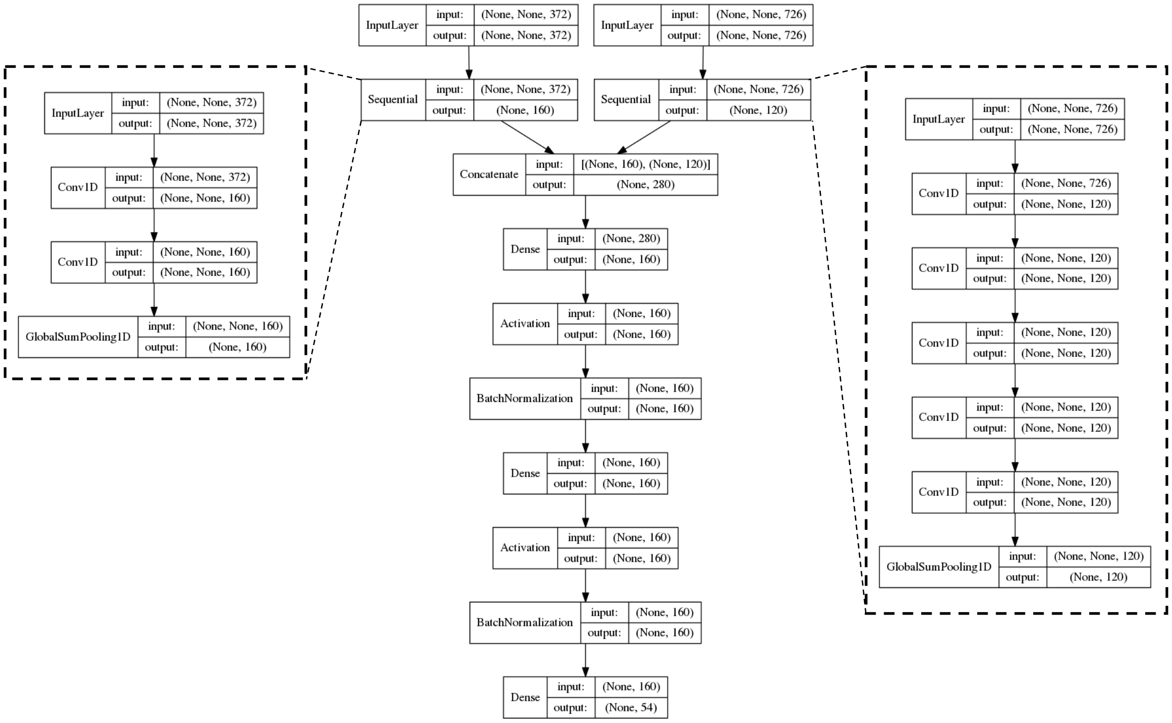


Transcript model

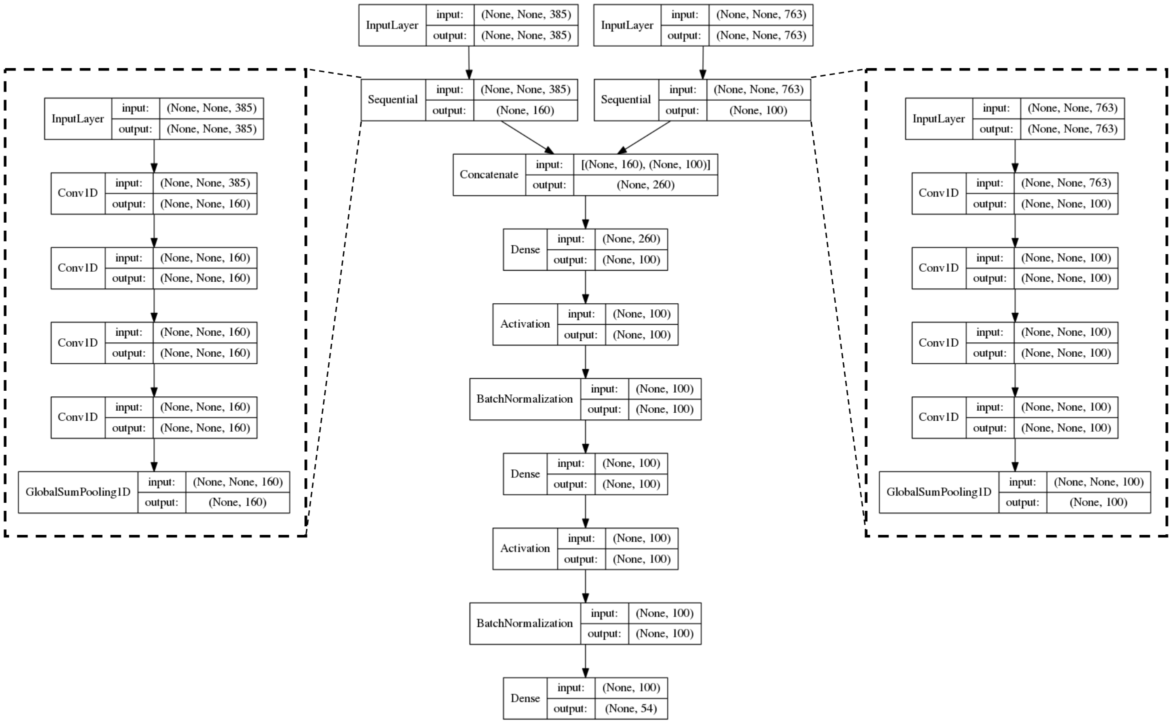


**Figure S17**. Structures of deep neural networks in DEcode.
