## Supplementary material for "Decoding differential gene expression": Figure S2

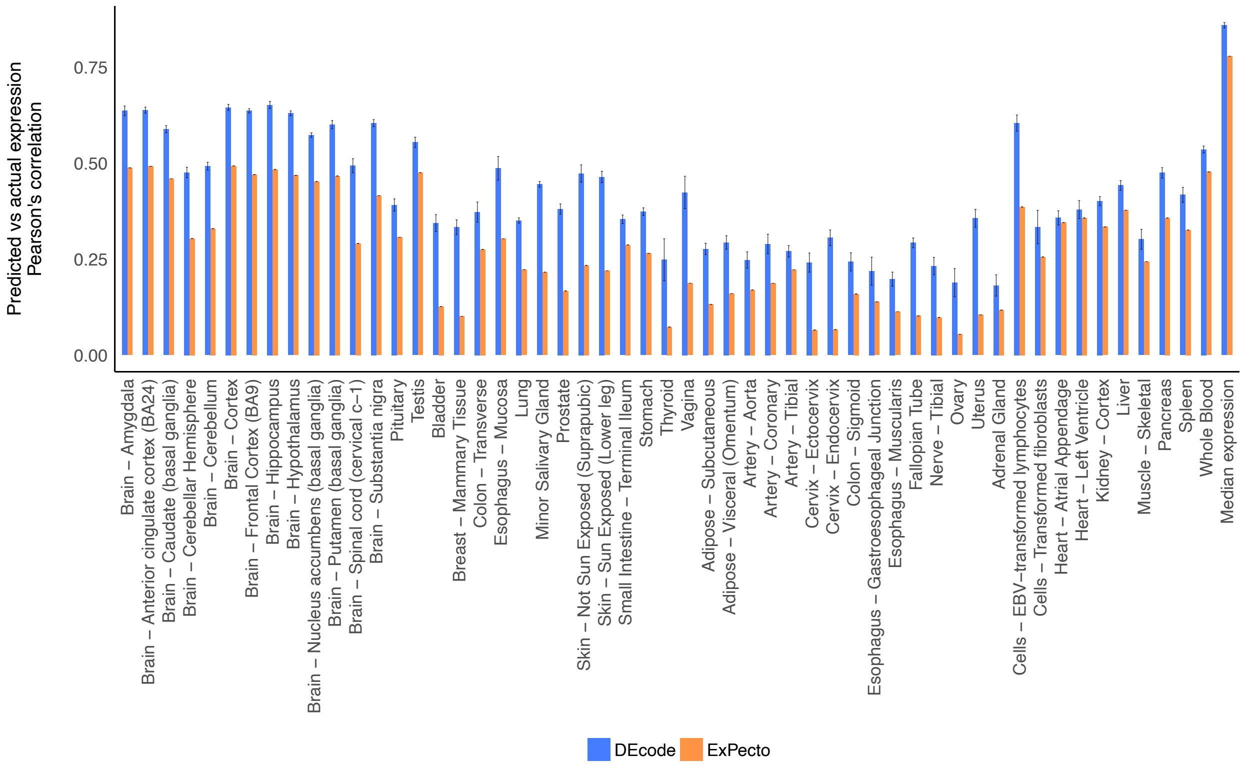


**Figure S2**. Performance comparison of DEcode with ExPecto with respect to correlation coefficient.
