## Supplementary material for "Decoding differential gene expression": Figure S3

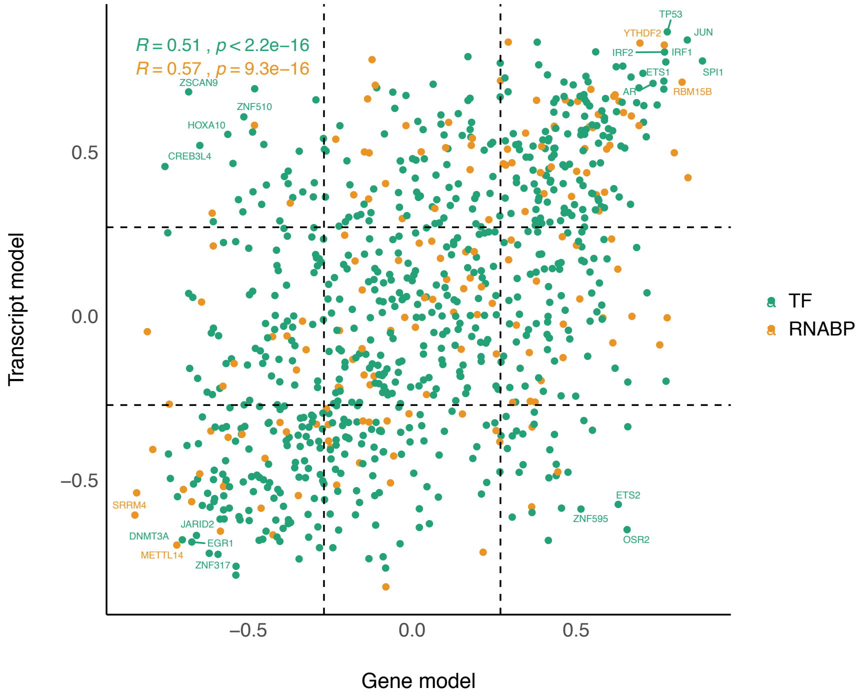


**Figure S3**. Comparison DeepLIFT scores between the gene-based model and the transcript-based model.
