## Supplementary material for "Decoding differential gene expression": Figure S4

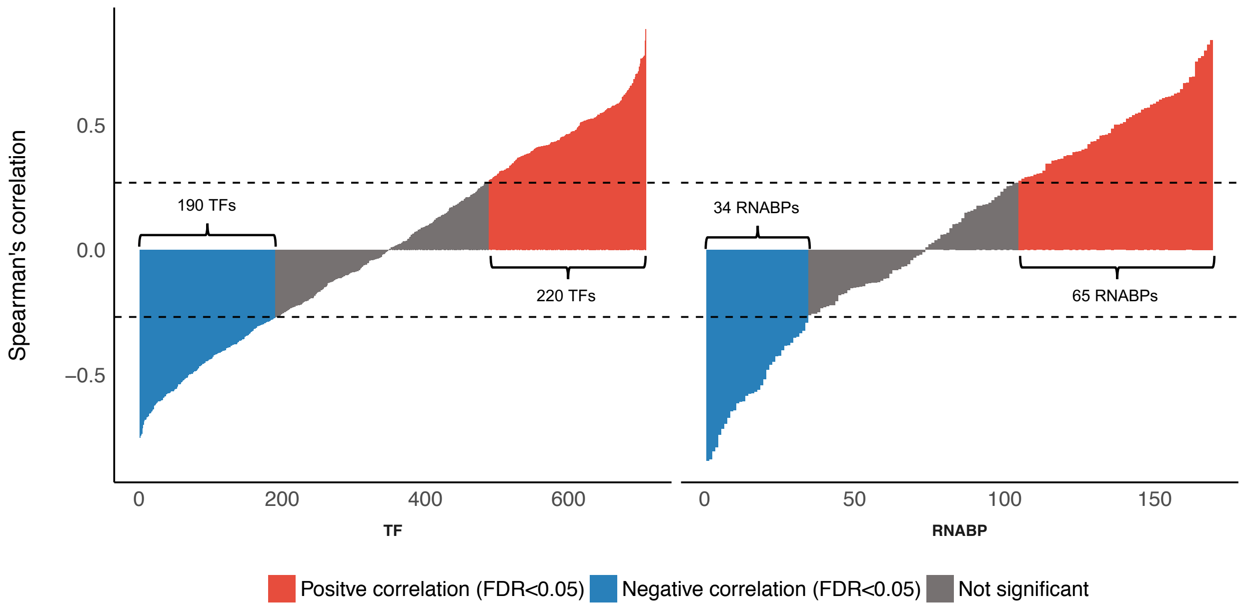


**Figure S4**. Correlation between DeepLIFT scores vs log2-TPMs for each of TFs and RNABPs. Spearman's correlation was used to evaluate the relation between DeepLIFT scores vs log2-TPMs. The Benjamini–Hochberg procedure was used to control the false discovery rate at 5%.
