## Supplementary material for "Decoding differential gene expression": Figure S5

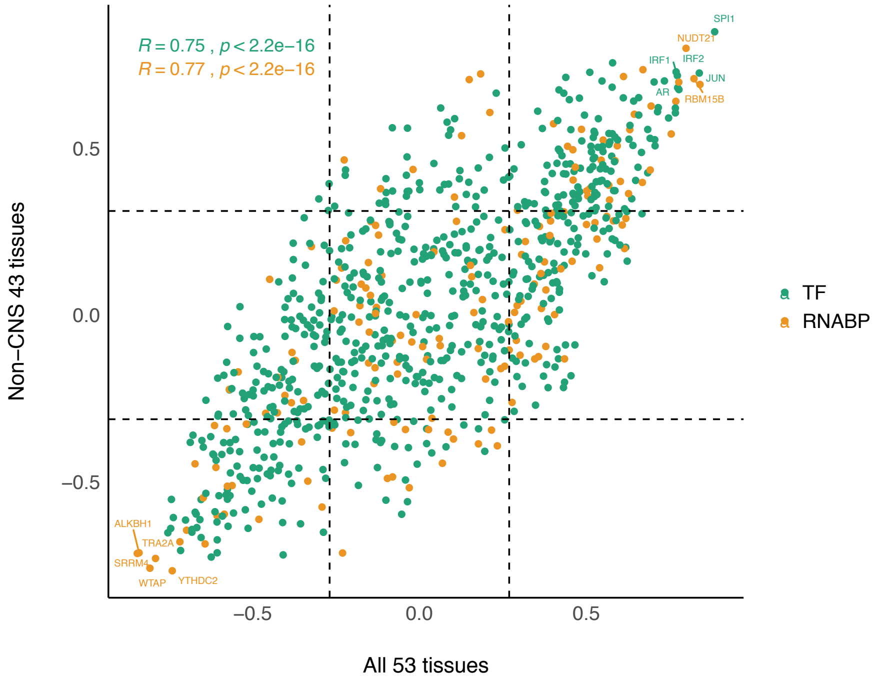


**Figure S5**. Relationships of regulators' DeepLIFT scores with their log2-TPMs with or without brain tissues. Spearman's correlations between DeepLIFT scores vs log2-TPMs for each of TFs and RNABPs were computed using all 53 tissues and 43 tissues without brain tissues, respectively. The Spearman's correlations from the two sets of tissues were contrasted by a scatter plot.
