## Supplementary material for "Decoding differential gene expression": Figure S6

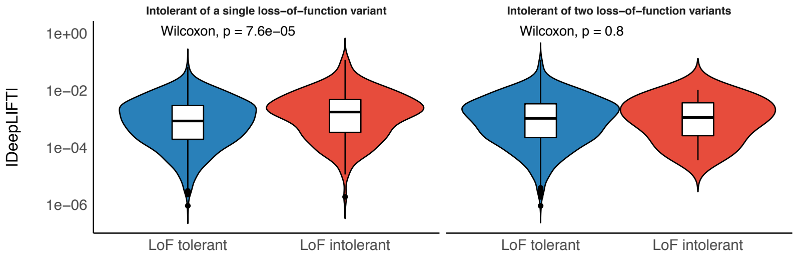

**Figure S6**. The overlap between the key regulators for the median absolute expression levels and LoF intolerant genes. The LoF intolerant genes were split into genes intolerant to heterozygous LoF mutations and genes intolerant to homozygous LoF mutations.
