## Supplementary material for "Decoding differential gene expression": Figure S7

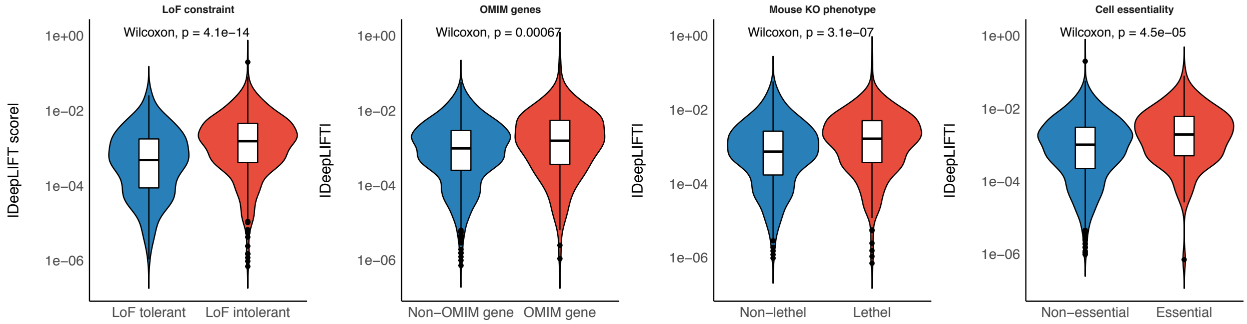


**Figure S7**. The overlap between the key regulators for the median absolute expression levels in the transcript-based model and external functional gene sets.
