## Supplementary material for "Decoding differential gene expression": Figure S8

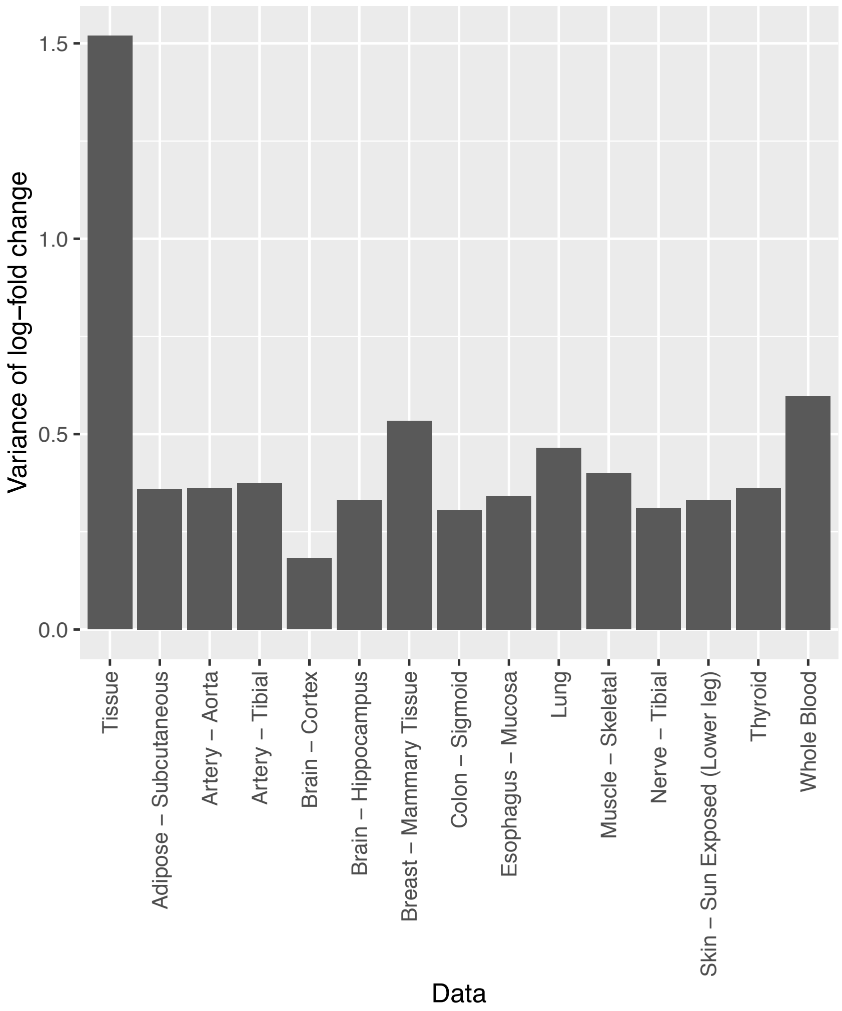


**Figure S8**. The variance of gene expression between tissues and within tissues. We computed variances in log-fold changes based on mean TPMs across 53 tissues and those based on TPMs across individuals in each tissue.
