## Supplementary material for "Decoding differential gene expression": Figure S10

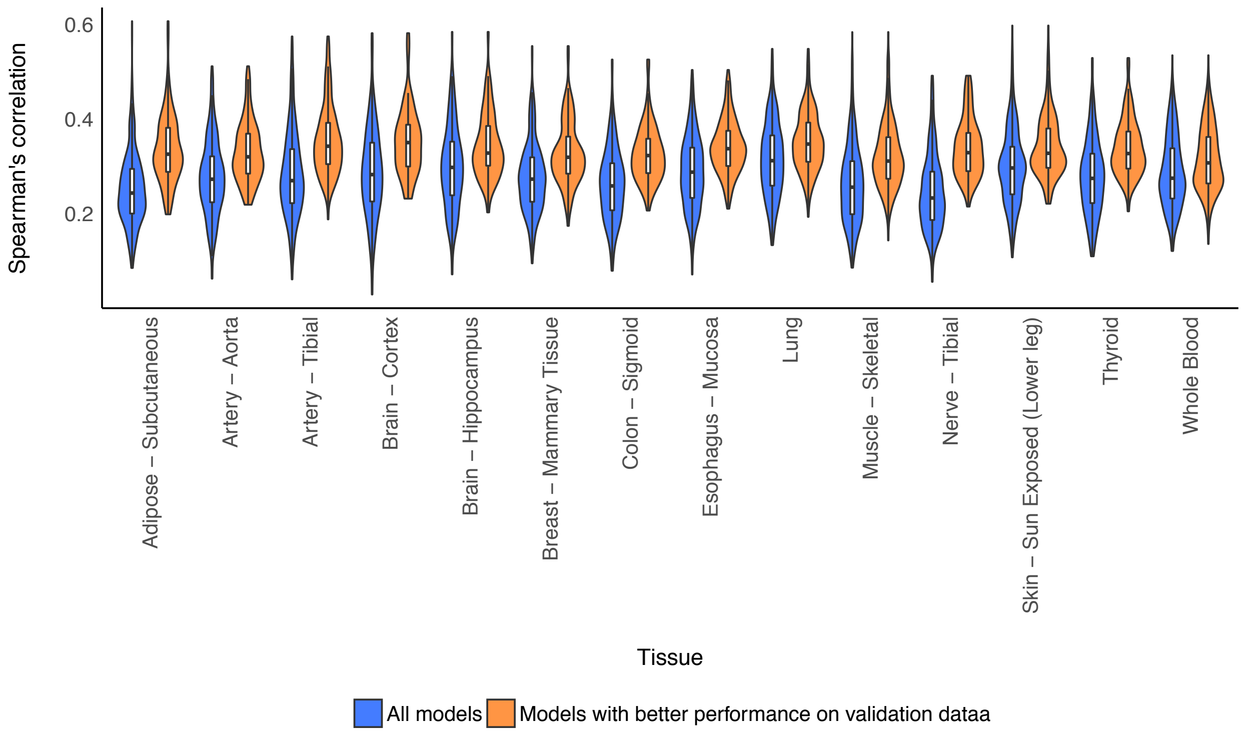


**Figure S10**. The predictive performances of the person-specific models. We computed Spearman's correlation between the predicted gene expression and the actual gene expression for each individual. We filtered out person-specific predictions from the models whose performances on validation data were less than 50% percentile of all individuals in each tissue.
