## Supplementary material for "Decoding differential gene expression": Figure S11

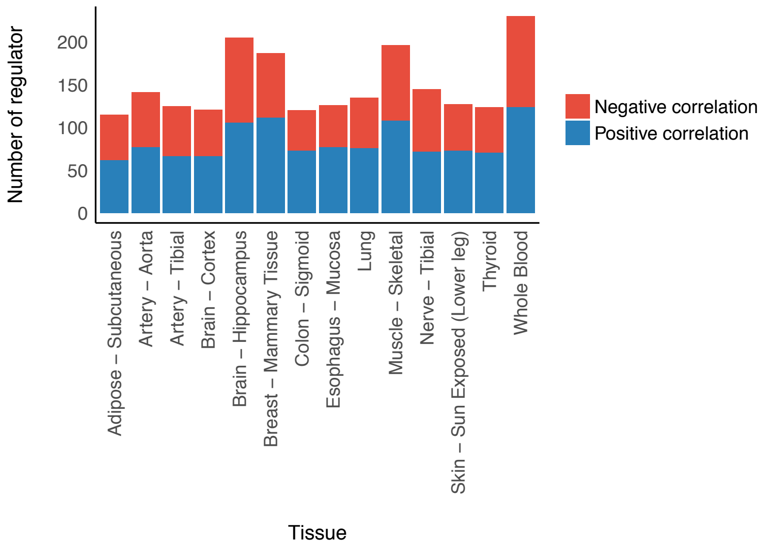


**Figure S11**. The number of regulators that showed significant relations between their DeepLIFT scores and log2-TPMs. Spearman's correlation was used to evaluate the relation between DeepLIFT scores vs log2-TPMs in each tissue. The absolute Spearman's correlation greater than 0.3 and the false discovery rate under 5% was defined as significant.
