## Supplementary material for "Decoding differential gene expression": Figure S12

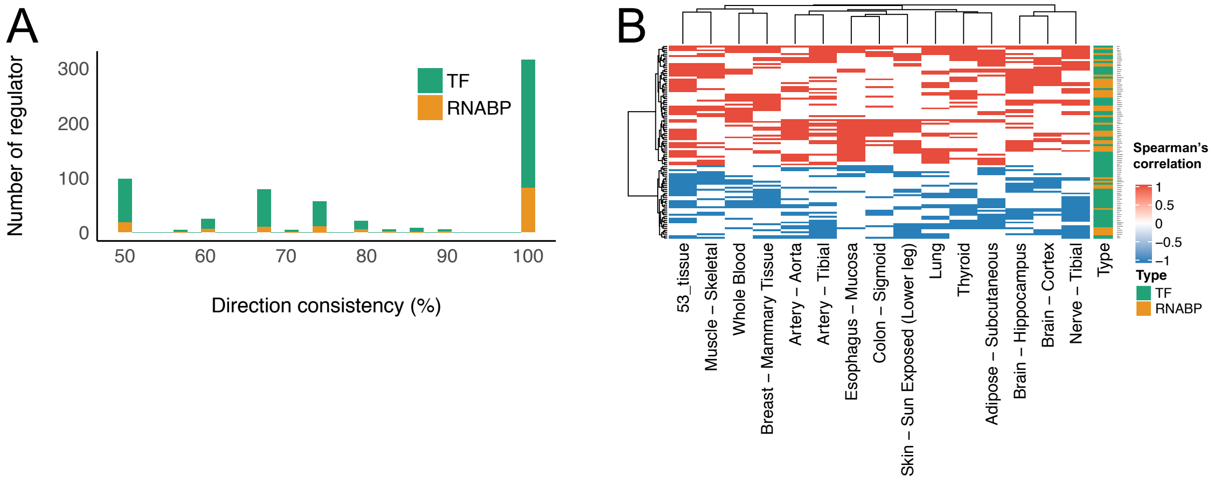


**Figure S12**. Consistency of the relationships of regulators' DeepLIFT scores with their log2-TPMs. (A) A histogram showing consistency of the relationships of regulators' DeepLIFT scores with their log2-TPMs. We selected the regulators that showed the significant relationships between their DeepLIFT scores with log2-TPMs in more than one tissue. Then, the consistency of directions of the correlations was computed and visualized as a histogram. (B) A heatmap showing the actual correlation between DeepLIFT scores and log2-TPMs. We selected 99 regulators that showed consistent relationships in more than four tissues.
