## Supplementary material for "Decoding differential gene expression": Figure S13

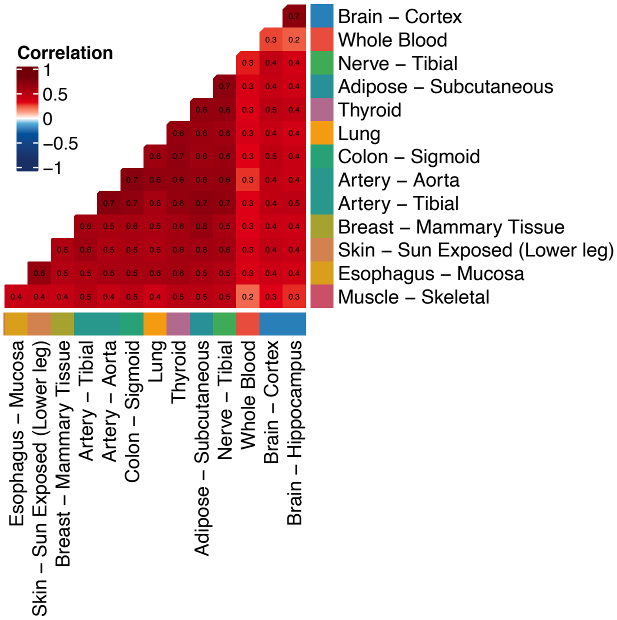


**Figure S13**. The heatmap shows the pairwise similarity of per-gene prediction accuracy between tissues. Spearman's correlation was used to evaluate the similarity and indicated in each element of the heatmap.
