## Supplementary material for "Decoding differential gene expression": Figure S14

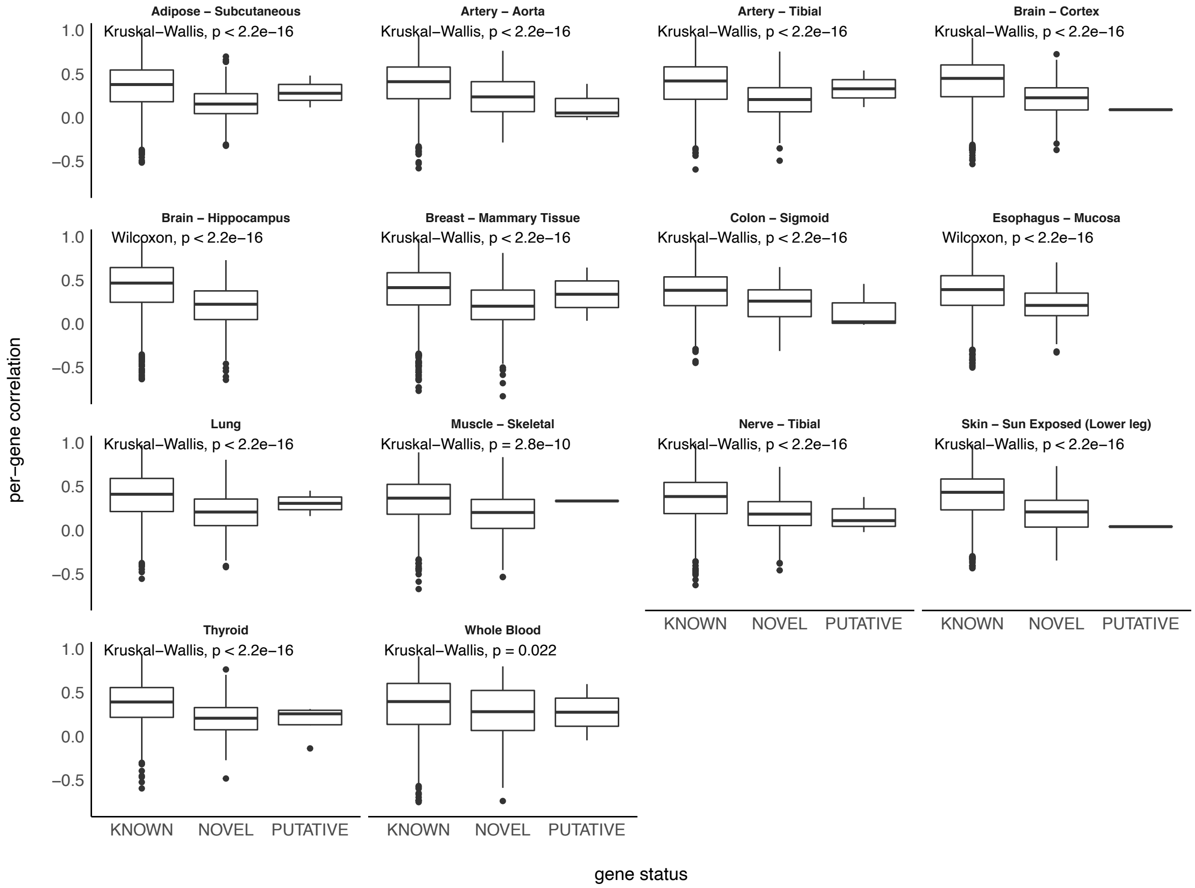


**Figure S14**. The relations of per-gene prediction accuracy with the gene status. The gene status was retrieved from the gtf file download from the GTEX portal. The gene status was assigned by the GENCODE consortium, representing genes are registered in multiple databases (KNOWN) or only in the GENCODE database (NOVEL or PUTATIVE).
