## Supplementary material for "Decoding differential gene expression": Figure S15

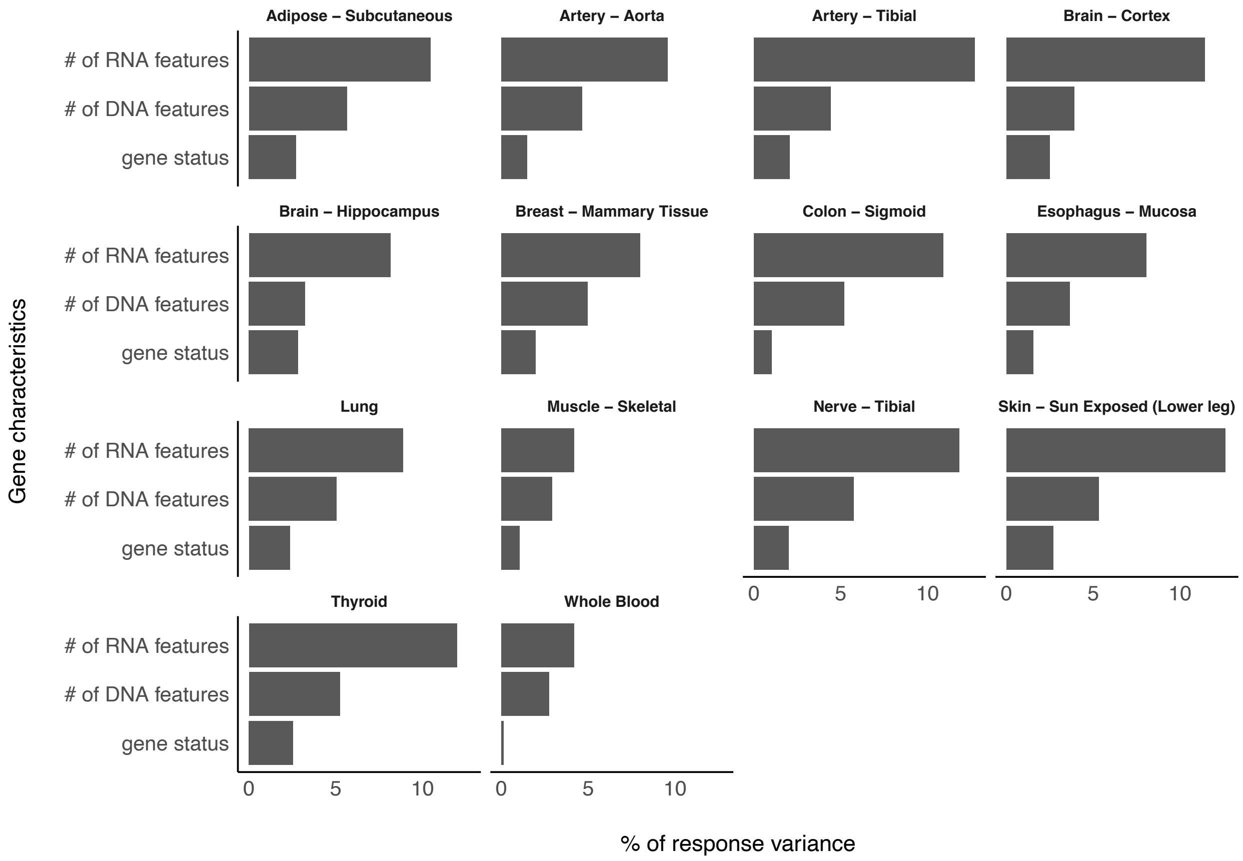


**Figure S15**. Decomposition of variance in the per-gene prediction accuracy explained by the gene status and the number of predictive features. To compare the variances in the per-gene prediction accuracy explained by the gene status and the number of DNA features and RNA features, we used a variance decomposition method called lmg that is implemented in relaimpo R package.
