## Supplementary material for "Decoding differential gene expression": Figure S16

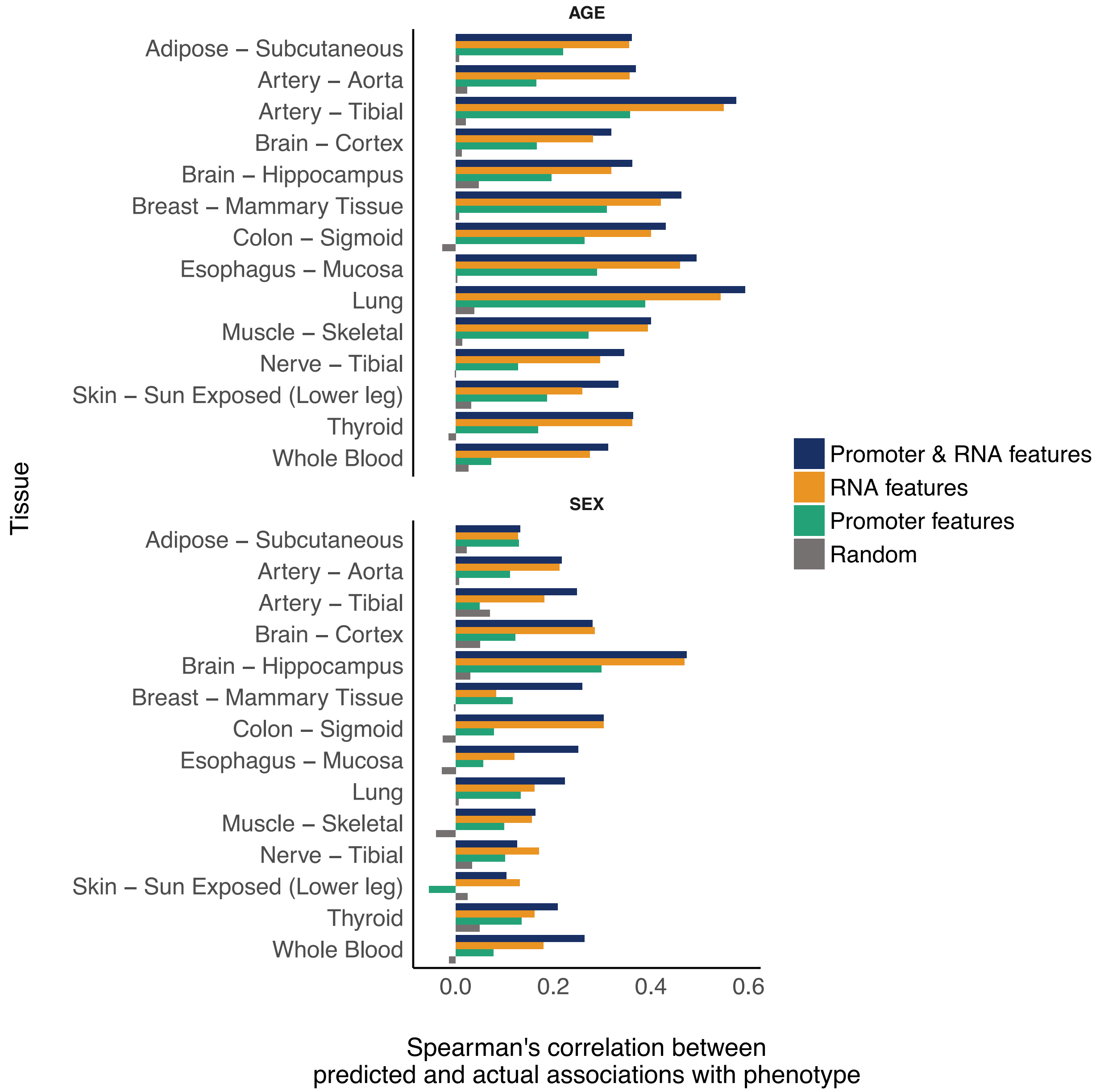


**Figure S16**. Comparison of performances in predicting phenotype-associated DEs with a distinct feature set. The bar plots show Spearman's correlation between t-statistics of DE using the predicted and the actual gene expression.
